# Distinct roles of H3K27me3 and H3K36me3 in vernalization response, maintenance and resetting in winter wheat

**DOI:** 10.1101/2023.12.19.572364

**Authors:** Xuemei Liu, Xuelei Lin, Min Deng, Bingxin Shi, Jinchao Chen, Haoran Li, Shujuan Xu, Xiaomin Bie, Xiansheng Zhang, Kang Chong, Jun Xiao

## Abstract

Winter plants rely on vernalization, a vital process for adapting to cold and ensuring successful reproduction. However, understanding the role of histone modifications in guiding the vernalization process in winter wheat is limited. In this study, we investigate the transcriptome and chromatin dynamics in the shoot apex throughout the life cycle of winter wheat in the field. Two core histone modifications, H3K27me3 and H3K36me3, exhibit opposite pattern on the key vernalization gene *VERNALIZATION1* (*VRN1*), correlated with its induction during cold exposure. Additionally, H3K36me3 remains high at *VRN1* after cold exposure, maintaining its active state. Mutations in FERTILIZATION-INDEPENDENT ENDOSPERM (TaFIE) and SET DOMAIN GROUP 8 (TaSDG8), writer complex components of H3K27me3 and H3K36me3, respectively, affect flowering time. Interestingly, *VRN1* loses its high expression after cold exposure memory in the absence of H3K36me3. During embryo development, *VRN1* is silenced with the removal of H3K36me3 in both winter and spring alleles. H3K27me3 is selectively added to the winter allele, influencing the cold exposure requirement for the next generation. Integrating gene expression with H3K27me3 and H3K36me3 patterns identified potential regulators of flowering. This study reveals distinct roles of H3K27me3 and H3K36me3 in controlling vernalization response, maintenance, and resetting in winter wheat.

**Significance Statement:** Vernalization, initially observed in cereals, lacks a comprehensive understanding of its underlying mechanism, particularly regarding chromatin-mediated transcriptional regulation in winter wheat. By delving into the transcriptome and chromatin dynamics in the shoot apex throughout winter wheat’s life cycle, we pinpointed two crucial histone modifications, H3K27me3 and H3K36me3, each playing distinct roles at different vernalization stages. H3K27me3 is implicated in establishing and resetting the extended cold exposure requirement for winter wheat, gradually diminishing during vernalization. On the other hand, H3K36me3 is crucial for maintaining *VRN1*’s active state post-cold exposure, contributing to the memory of the vernalization treatment. Additionally, the integration of transcriptome and histone modification profiles unveiled potential novel regulators of flowering in winter wheat.

## Introduction

Winter wheat undergoes vernalization, a process involving prolonged exposure to low temperatures, to shift from vegetative to reproductive growth and enhance grain production (1). This crucial process ensures proper flowering timing and maximizes grain yields (2–5). The regulation of vernalization in winter wheat is intricate, involving a network of regulatory genes, with *VRN1*, *VRN2*, and *VRN3* being extensively studied (6–8). The interactions among these genes play a central role in controlling the vernalization response and requirement (9, 10). The activation of *VRN1* is pivotal for initiating the floral transition, and the polymorphism at the *VRN1* locus determines the need and duration of cold exposure (11–14). Research has revealed that *VRN1* expression is subject to regulation at both transcriptional (15, 16) and post-transcriptional levels (17), as well as through epigenetic modifications (17–20).

Chilling temperatures have been demonstrated to induce histone modifications that facilitate the activation of *VRN1*. Specifically, the removal of tri-methylation from lysine 27 on histone H3 (H3K27me3) at the *VRN1* locus enables its activation. Additionally, histone marks associated with active transcription, such as H3K4me3, increase at the promoter and first intron regions of *VRN1*, promoting gene expression (17, 18). Beyond H3K27me3 and H3K4me3, H3K36me3 plays a crucial role in reversing H3K27me3 at the FLOWERING LOCUS C (*FLC*) locus in *Arabidopsis* during the vernalization process (21, 22). After cold exposure, the transcriptional status of key genes is maintained mitotically until spring when the flowering transition occurs (2, 23). In *Arabidopsis*, the silenced status of *FLC* is preserved through the spreading of H3K27me3 across the entire gene body after returning to warm temperatures (24). In barley and wheat, winter-activated *VRN1* expression remains high post-cold exposure (4), but the connection to active histone modifications and the specific histone modification involved is yet to be studied.

Crucially, the epigenetic status of key regulators in flowering regulation in response to vernalization must be reset to maintain the vernalization requirement for the next generation (25–27). In *Arabidopsis*, the coupling of H3K27me3 to *FLC* silence persists in both male and female gametes after vernalization (28, 29). The repressed state of *FLC* is maternally transmitted to early embryos, where the reactivation of *FLC* occurs with the function of the pioneer transcription factor (TF) *LEAFY COTYLEDON 1* (*LEC1*) and other B3-domain TFs, including *LEC2*, *FUSCA3* (*FUS3*), and *ABSCISIC ACID INSENSITIVE* (*ABI3*) (30–32). Notably, active demethylation of H3K27me3 by histone demethylase EARLY FLOWERING 6 (ELF6) is also necessary during this process (25). For wheat, the timing of when active *VRN1* is reset and the factors involved remain largely unknown.

In addition to *VRN1*, other candidates are proposed to play a role in the regulation of the vernalization response in wheat, based on their expression patterns. These include potential *FLC* orthologs *ODDSOC2* (*TaOS2*) and *TaOS1*, although their impact on flowering time appears to be marginal, as indicated by genetic data from mutations (33, 34). Furthermore, a global analysis of the H3K27me3 and H3K4me3 patterns in *Brachypodium* at different vernalization stages (V0, V30, V30N) has identified additional loci involved in mediating the epigenetic memory of flowering competency acquired during the vernalization process (35). Thus, both transcriptional and histone modification patterns may serve as indicators for the identification of potential regulators for vernalization-promoted flowering.

In this study, we delved into the role of the chromatin landscape in driving the vernalization response, memory, and resetting in winter wheat by profiling chromatin accessibility, core histone modifications, and the histone variant H2A.Z in the shoot apex throughout an entire growth season in the field. We observed antagonistic modifications of H3K27me3 and H3K36me3 at the *VRN1* locus during the entire life cycle of winter wheat. Through assessing flowering time, *VRN1* transcription levels, and relative histone modifications in genome-edited Polycomb Repressive Complexes 2 (PRC2) and SDG8 mutants, we unveiled distinct functions of H3K27me3 and H3K36me3 in mediating the vernalization response, memory, and resetting processes. Moreover, integrating transcriptome patterns with the profiles of H3K27me3 and H3K36me3 allowed the identification of additional *VRN1*- and *FLC*-pattern-like genes for potential regulation of flowering in wheat.

## Results

### Profiling the transcriptome and chromatin landscape in the shoot apex across the life-cycle of winter wheat in field

Winter wheat is traditionally sown in early October and experiences a prolonged cold winter from November to early March, with temperatures ranging from -10°C to 5°C in Beijing, China (Fig. 1A). As temperatures rise above 10°C in mid-March, wheat jointing stage occurs. We conducted a comprehensive study on the transcriptome and chromatin dynamics during the vernalization response, maintenance, and resetting stages of winter wheat cultivar Kenong 9204 (KN9204), using samples of shoot apex and developing seed/embryo from field-grown wheat (Fig. 1A). Our efforts resulted in the generation of 21 RNA-seq libraries, 14 ATAC-seq libraries, and 94 CUT&Tag libraries, targeting various histone modifications and variant (H3K27ac, H3K27me3, H3K36me3, H3K4me3, and H2A.Z) (Fig. 1B). Quality assessments revealed excellent reproducibility for both transcriptome and epigenome data (Fig. S1A, D, E), and the distribution of histone modifications and chromatin accessibility profiles across the genome and genes aligned with their known roles in transcriptional regulation, as previously reported (36, 37) (Fig. S1B, C).

**Fig. 1.**
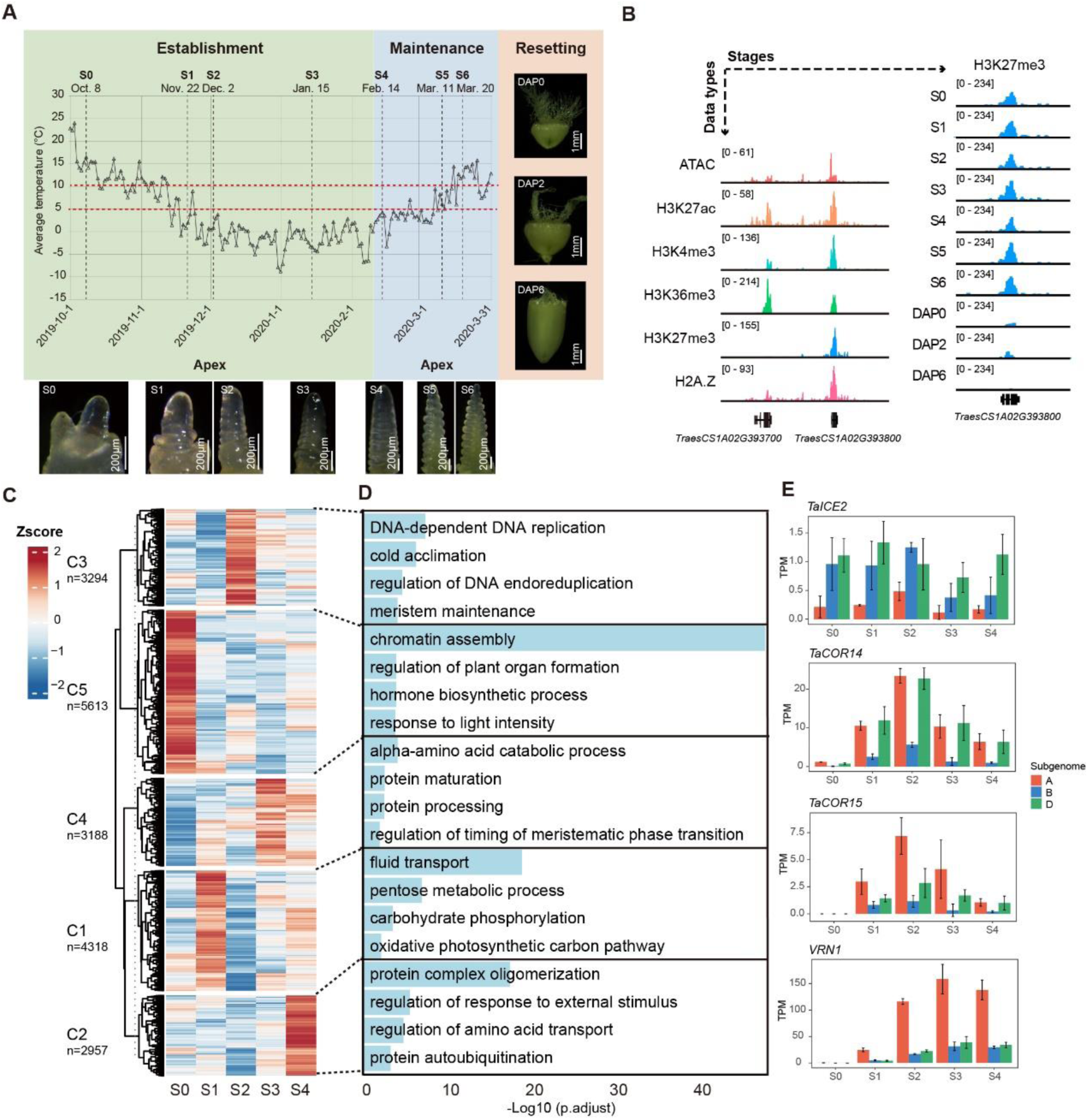
Profiling the transcriptome and epigenome in the shoot apex during vernalization in winter wheat. **A.** Temperature at different sampling time points. DAP: Days After Pollination. **B.** Data types and sampling time points in this study. **C.** K-means clustering heatmap of DEGs. The color represents the level of transcription. **D.** GO enrichment analysis of DEGs in all clusters. **E.** Transcription patterns of *TaICE2*, *TaCOR14*, *TaCOR15* and *VRN1* during vernalization.

Differential expression analysis was conducted for S0 (pre-vernalization), S1 and S2 (during vernalization), and S3 and S4 (fully vernalized), revealing 19,370 differentially expressed genes (DEGs) (Table S1). Subsequent K-means clustering classified DEGs into five clusters with distinct expression patterns (Fig. 1C). Notably, genes in cluster C5 exhibited high expression before cold exposure, while genes in clusters C1 and C3 were highly induced during cold treatment, and those in clusters C4 and C2 were predominantly expressed in the fully vernalized state. Gene Ontology (GO) enrichment analysis shed light on the functions of stage-specific highly expressed genes (Fig. 1D). Before vernalization, wheat developments fast, genes were associated with chromatin assembly, plant organ formation, hormone biosynthetic, and response to light intensity (C5). During vernalization, winter wheat initially acquired cold tolerance, with genes related to fluid transport, pentose metabolic, carbohydrate phosphorylation, oxidative photosynthetic carbon pathway are highly expressed in the S1 stage (C1), and cold acclimation genes upregulated in the S2 stage (C3). Notably, key cold-responsive modules, including the *INDUCER OF CBF EXPRESSION-C REPEAT BINDING FACTOR-COLD RESPONSIVE* (*ICE-CBF-COR*) cluster (38), were highly expressed at the S2 stage (Fig. 1E), indicating early cold tolerance acquisition. In the vernalization maturation stage (39), genes related to amino acid catabolic, protein maturation, protein complex oligomerization, and amino acid transport were highly expressed (C4, C2), including the crucial flowering time regulator, *VRN1* (6) (Fig. 1E). While both *VRN1* and *TaCOR14/15* responded to low temperatures, their expression patterns differed (Fig. 1E). *TaCOR14/15* exhibited high expression levels only during early cold exposure, followed by a decline, whereas *VRN1*’s expression increased gradually with prolonged cold exposure until the vernalization effect was fully established (Fig. 1E). These distinct expression patterns corresponded to their functions in cold tolerance and the measurement of cold duration for flowering promotion (3). In summary, the transcriptome effectively captures the various stages of vernalization in winter wheat, highlighting the differentiation between short-duration cold tolerance genes and long-duration vernalization genes.

### Dynamics of chromatin states during vernalization establishment and maintenance

We utilized chromHMM (40) to explore the dynamic alterations in chromatin states throughout vernalization, integrating multiple histone modifications and variant data. Models with 5 to 25 states were trained, and 11 chromatin states were selected based on their similarities (Fig. S2A). Each chromatin state was assigned a descriptive label according to enriched histone modifications, distance to genes, and expression levels of nearby genes (Fig. 2A). These 11 states were grouped into four based on transcriptional activity (Fig. 2A). Group G1, marked by high gene expression, included states E1, E2, E3, and E4, characterized by the presence of H3K36me3, associated with active transcription (Fig. S2B). In contrast, G3, featuring low gene expression, comprised states E8, E9, and E10, enriched in H3K27me3, a repressive modification inversely correlated with transcription (Fig. S2B). E8 displayed a bivalent chromatin state (41), showing both H3K27me3 and active histone marks H3K4me3 and H3K27ac. E4, rich in H3K27ac, H3K4me3, and H2A.Z, exhibited the highest chromatin accessibility, while E1, E2, and E3, with high transcription levels, displayed lower accessibility (Fig. 2A), aligning with the idea that transcriptional regulatory regions are generally more accessible than gene bodies (42).

**Fig. 2.**
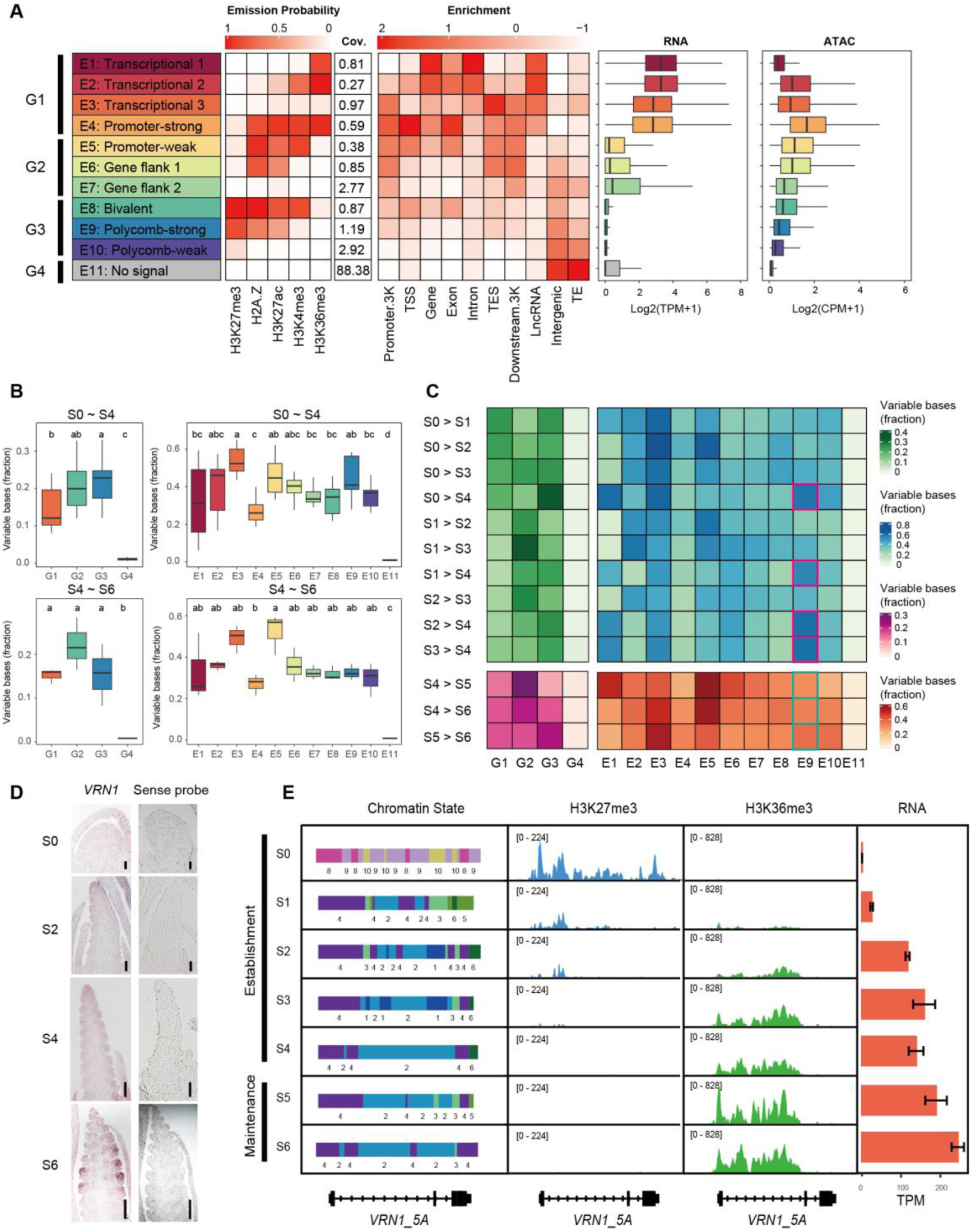
The chromatin state changes during vernalization establishment and maintenance. **A.** 11 chromatin states informed from all CUT&Tag data. Each row represents a chromatin state and each column respectively represents histone marker enrichment (Emission Probability), genomic coverage, genomic location distribution (Enrichment), gene expression level and chromatin accessibility level. **B.** Comparison of variable bases fraction between different chromatin state. The letters represent the results of multiple comparisons, with a significance level of 0.05. **C.** The fraction of bases that change chromatin state between different stages. **D.** In situ hybridization of *VRN1* at S0, S2, S4, and S6 stages. **E.** The chromatin state, H3K27me3, H3K36me3 and transcriptional level changes of *VRN1_5A* from S0 to S6.

Exploring the transition of chromatin states during vernalization establishment (S0-S4) revealed more significant changes in G2 and G3 compared to G1 and G4 (Fig. 2B). E3, E5, and E9 exhibited the most substantial changes (Fig. 2B), with E9 in the fully vernalized S4 stage displaying a distinct profile (Fig. 2C). Most regions in E9 state at S0, S1, S2, S3 remained in E9 state at S4 (Fig. S2C). However, genes with chromatin state changed at S4, especially transitioning into E1 or E2 state, were mainly associated with flowering development, auxin signaling, and flower organ specification (Fig. S2D), indicating a crucial role for H3K27me3 in vernalization establishment. After extended exposure to winter cold, wheat plants initiated bolting in late March, relying on the ‘memory’ of the vernalized state (3). Notably, E9 state remained more stable from S4 to S5 and S6 (Fig. 2B, C), suggesting that H3K27me3 may not be involved in maintaining vernalization memory.

Further examination of the core regulator, VRN1, revealed quantitative induction in response to cold exposure from S0 to S4, maintaining high expression in S6 after cold treatment in the shoot apex (Fig. 2D). This pattern aligns with its role in promoting flowering and regulating spikelet development (6, 43). Before vernalization, *VRN1* exhibited minimal expression, with its promoter and first intron (*TraesCS5A02G391700*) covered by H3K27me3 within chromatin state E9 (Fig. 2E). Continuous exposure to low temperatures led to a gradual decrease in H3K27me3 levels at *VRN1*, accompanied by a progressive increase of H3K36me3 levels, resulting in a shift from E9 to E2 and E4 chromatin states and continuous upregulation of transcription (Fig. 2E, Fig. S2E, F). A similar inverse pattern of H3K27me3 and H3K36me3 was observed in the core vernalization gene *FLC* in *Arabidopsis thaliana* (22). Moreover, upon return to normal temperatures after vernalization, H3K27me3 enveloped the entire *FLC* locus to inhibit transcription in *Arabidopsis* (21). Similarly, in wheat, upon transitioning to normal temperatures (S5 and S6 stages) after vernalization, the presence of H3K36me3 coincided with high *VRN1* expression (Fig. 2E, Fig. S2E, F). These findings highlight the changing chromatin states during vernalization in winter wheat, emphasizing the importance of H3K27me3 and H3K36me3 in regulating key flowering genes like *VRN1*, with the conserved roles across various plant species.

### Modulation of H3K27me3 or H3K36me3 affects vernalization-induced flowering in winter wheat

Giving the potential pivotal role of H3K27me3 and H3K36me3 in orchestrating the vernalization response, we conducted a genetic analysis to investigate their influence on flowering by manipulating these histone modifications. Through CRISPR-Cas9 genome editing, we generated mutations targeting the “writer” complex responsible for H3K27me3 and H3K36me3 in the winter wheat KN199 (Fig. S3A, B), a derived cultivar from KN9204 with moderate genetic transformation efficiency (44). Knocking out all three subgenomes of *TaFIE*, a core component of the PRC2 (45), proved embryonic lethal (37). Therefore, we opted for a mild mutation, the *Tafie-cr-87* line, which exhibited a frameshift mutation in the A and D subgenomes and two amino acid deletions in the B subgenome of *TaFIE* (Fig. S3A) (37). Notably, the global H3K27me3 level was reduced in *Tafie-cr-87* compared to KN199 at shoot apex (Fig. S3C).

We assessed the flowering times of *Tafie-cr-87* and KN199 under various vernalization treatments, including 0 day (V0), 20 days (V20), and 30 days (V30) of exposure to 4°C. Under V0 and V20 conditions, *Tafie-cr-87* displayed earlier flowering compared to KN199, while no differences were observed under V30 conditions with saturated cold exposure (Fig. 3A, B). Subsequently, we examined the expression levels of *VRN1* in *Tafie-cr-87* and KN199 under different vernalization treatments. *VRN1* expression was significantly elevated in *Tafie-cr-87* compared to KN199 at the seedling stage under V0 and V20 conditions, while no differences were observed under V30 conditions (Fig. 3C, Fig. S3E). Additionally, H3K27me3 levels were reduced at the *VRN1* promoter region in *Tafie-cr-87* compared to KN199 under V0 and V20 conditions, but H3K27me3 diminished in both plants under V30 conditions (Fig. 3D, Fig. S3F). These findings suggest that PRC2-H3K27me3 primarily inhibits *VRN1* expression without or under insufficient vernalization treatment, with the repressive effect diminishing with prolonged cold exposure.

**Fig. 3.**
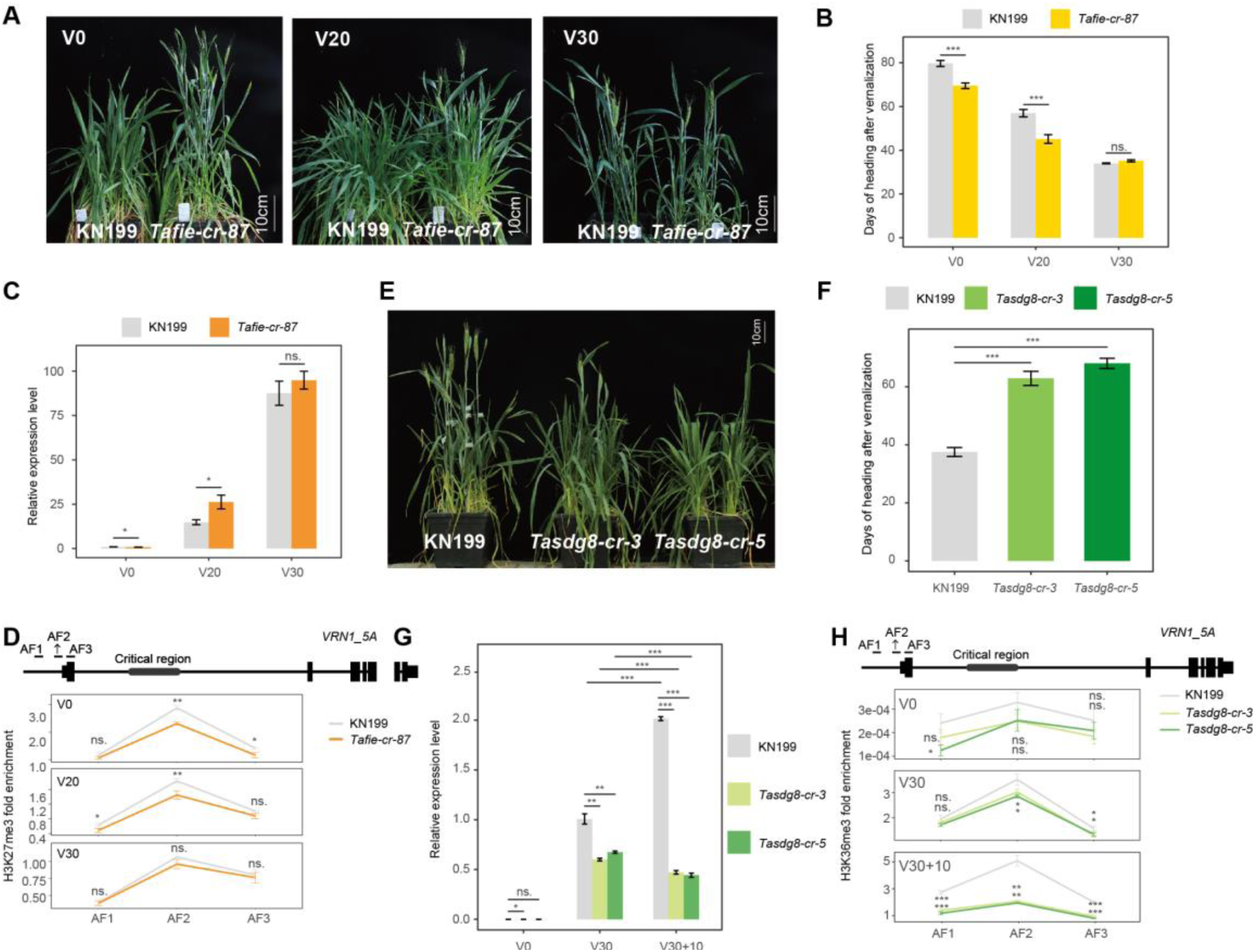
Mutation of TaFIE and TaSDG8 affects vernalisation-induced flowering. **A, B.** Photos (A) and heading time (B) of wild type (KN199) and *Tafie* crispr/cas9 line (*Tafie-cr-87*) under different vernalization treatments. V0, V20, and V30 represent 0, 20, and 30 days of treatment at 4°C respectively. **C**. The relative expression level of *VRN1_5A* in seedlings of wild type and *Tafie* crispr/cas9 line under different vernalization treatments. **D**. The H3K27me3 level at different location of *VRN1_5A* in seedlings of wild type (KN199) and *Tafie* crispr/cas9 line (*Tafie-cr-87*) at heading stage under different vernalization treatments. **E, F.** Photos (E) and heading time (F) of wild type (KN199) and *Tasdg8* crispr/cas9 lines (*Tasdg8-cr-3*, *Tasdg8-cr-5*) under 30 days of vernalization treatment at 4°C. **G.** The relative expression level of *VRN1_5A* in seedlings of wild type (KN199) and *Tasdg8* crispr/cas9 (*Tasdg8-cr-3*, *Tasdg8-cr-5*) lines under different vernalization treatments. V30+10 represents 30 day days of treatment at 4°C and subsequent 10 days treatment at normal temperature. **H**. The H3K36me3 level at different location of *VRN1_5A* in seedlings of wild type (KN199) and *Tasdg8* crispr/cas9 (*Tasdg8-cr-3*, *Tasdg8-cr-5*) lines under different vernalization treatments.

In *Arabidopsis*, SDG8 functions as a methyltransferase for H3K36me3 in regulating flowering (46). In wheat, three homologues of *SDG8* form a triads, with A-, B- and D- subgenome *TaSDG8* exhibiting high expression in the shoot apex (Fig. S3D). To investigate the role of H3K36me3 in vernalization-induced flowering, we generated two *TaSDG8-a/b/d* mutant lines using CRISPR-Cas9 with reduced global H3K36me3 level (Fig. S3B, C), and evaluated their flowering time with vernalization treatment. Both *Tasdg8-cr-3* and *Tasdg8-cr-5* lines exhibited delayed flowering (Fig. 3E, F). Assessment of *VRN1* expression levels (Fig. 3G, Fig. S3G) and H3K36me3 status (Fig. 3H, Fig. S3H) in KN199 and *Tasdg8-cr-3/5* lines under V30 and V30+10 (exposure to 4°C for 30 days and convert to normal temperature for 10 days) conditions revealed that *VRN1* levels decreased along with reduced H3K36me3 levels in *Tasdg8-cr-3/5* lines under V30+10 compared to V30 conditions. In contrast, increased expression and H3K36me3 level was observed in KN199 under these conditions (Fig. 3G, H). Thus, the accumulation of H3K36me3 in *VRN1*, induced by prolonged cold exposure, contributes to the memory of vernalization treatment post-cold exposure in winter wheat.

### Resetting H3K27me3 and H3K36me3 at *VRN1* during embryogenesis to restore vernalization requirement in winter wheat

Following flowering, the vernalization memory must be wiped clean in the offspring to prevent the inheritance of the vernalized state across generations (25, 27). In *Arabidopsis*, the resetting of the vernalization requirement takes place during embryogenesis and involves the reactivation of *FLC* (25, 30). This process is associated with the reset of epigenetic modifications, such as H3K27me3 at the *FLC* locus (27). Given that VRN1 is the pivotal locus determining the vernalization requirement in wheat, we examined the expression pattern of *VRN1* during embryogenesis using in situ hybridization in winter wheat KN9204 (Fig. 4A). Prior to fertilization, we observed a robust signal of *VRN1* transcript in the embryo sac. However, *VRN1* expression gradually decreased with increasing days after pollination (DAPs), becoming barely detectable at DAP6 in the developing embryo and entirely undetectable at DAP14 (Fig. 4A). This indicates that *VRN1* expression is repressed during embryogenesis, mirroring the resetting process of *FLC* in *Arabidopsis* but with an opposite trend.

**Fig. 4.**
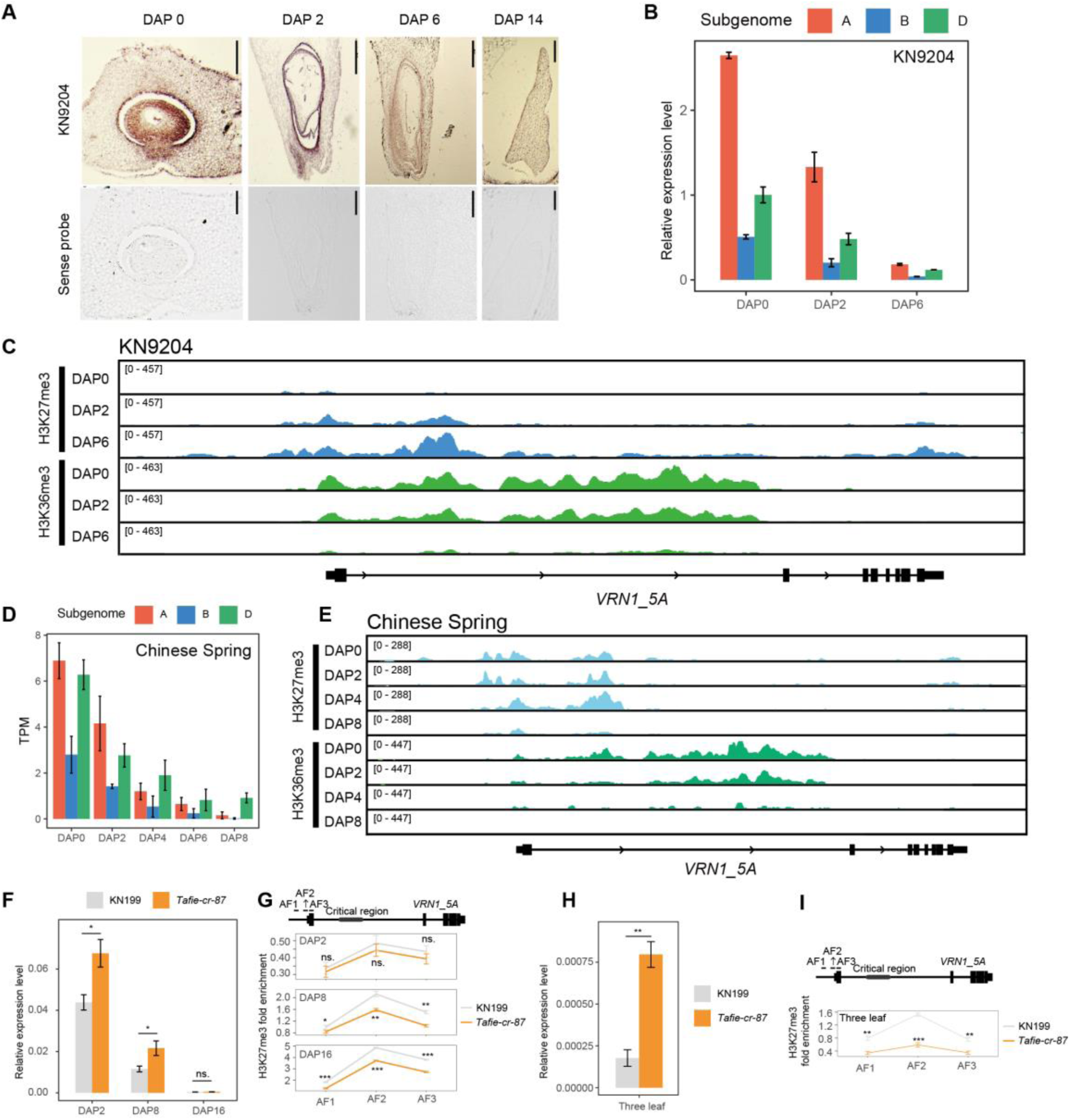
H3K27me3 restoration associates with the reset of vernalization requirement during embryogenesis. **A.** In situ hybridization of *VRN1* at DAP0, DAP2, DAP6, and DAP14 stages. **B.** The expression level of *VRN1* during embryogenesis in winter wheat KN9204. **C.** The H3K27me3 and H3K36me3 changes of *VRN1_5A* during embryogenesis in winter wheat KN9204. **D.** The expression level of *VRN1* during embryogenesis in spring wheat Chinese Spring. **E.** The H3K27me3 and H3K36me3 changes of recessive allele *VRN1_5A* during embryogenesis in spring wheat CS. **F.** The relative expression level of *VRN1_5A* during embryogenesis of wild type (KN199) and *Tafie* crispr/cas9 (*Tafie-cr-87*) line. **G.** The H3K27me3 level at different location of *VRN1_5A* during embryogenesis of wild type (KN199) and *Tafie* crispr/cas9 line (*Tafie-cr-87*). **H.** The relative expression level of *VRN1_5A* in leaves of wild type (KN199) and *Tafie* crispr/cas9 (*Tafie-cr-87*) line. **I.** The H3K27me3 level at different location of *VRN1_5A* in leaves of wild type (KN199) and *Tafie* crispr/cas9 line (*Tafie-cr-87*).

To delve deeper into the resetting of the vernalization “memory” in wheat, we utilized developing seeds at DAP0, DAP2, and DAP6 to conduct CUT&Tag analysis for four histone modifications (H3K27ac, H3K27me3, H3K36me3, H3K4me3) (Fig. 1A, B). Consistent with *in situ* hybridization findings, we validated a transcriptional decline of *VRN1* during embryogenesis (Fig. 4B). Particularly noteworthy was the shift in the chromatin state at the *VRN1-5A* locus from an active to a repressed state during embryogenesis (Fig. 4C). Notably, H3K36me3 levels significantly decreased at proximal promoter and within the *VRN1-5A* gene body, especially in the first intron region, from DAP0 to DAP6. Conversely, the repressive histone mark H3K27me3 gradually increased, encompassing the entire genic region, proximal promoter, and 3’UTR regions of *VRN1-5A* at DAP6 (Fig. 4C). Though there are various among different subgenomes of *VRN1* triads, the general pattern is similar (Fig. S4A). This underscores the significance of the reciprocal patterns of H3K27me3 and H3K36me3 in the transcriptional resetting of *VRN1* during embryogenesis. Additionally, we identified a similar pattern of reduced *VRN1* expression in spring wheat Chinese Spring (CS) for both the recessive *vrn1-5A vrn1-5B* and dominant *Vrn-5D* alleles from DAP0 to DAP8 (Fig. 4D) (37). Coinciding with reduced transcription, H3K36me3 levels declined in all three sub-genomes (Fig. 4E, Fig. S4B). However, we did not observe the accumulation of H3K27me3 during embryogenesis at DAP8 (Fig. 4E, Fig. S4B). This difference is likely linked to the variation that *VRN1* can be detected at the seedling stage, especially three-leaf stage, in CS but merely in KN9204 (Fig. S4C).

To ascertain whether the restoration of H3K27me3 at the *VRN1* locus during embryogenesis contributes to the repression of *VRN1* prior to vernalization, we compared mRNA levels and H3K27me3 statuses at the *VRN1* locus in KN199 and *Tafie-cr-87* during embryogenesis (DAP2, DAP8, DAP16) and the three-leaf seedling stages before cold exposure. Both KN199 and *Tafie-cr-87* exhibited a decline in *VRN1* mRNA levels during embryogenesis, with slightly or no significant difference between the two genotypes at DAP16 (Fig. 4F, Fig. S4D, F). However, H3K27me3 accumulated in KN199 but remained less enriched in *Tafie-cr-87* at 8 and DAP16 (Fig. 4G, Fig. S4E, G). At the three-leaf seedling stage, *Tafie-cr-87* displayed higher *VRN1* expression levels than KN199 (Fig. 4H, Fig. S4H, J), accompanied by lower H3K27me3 levels (Fig. 4I, Fig. S4I, K). In conclusion, the resetting of H3K27me3 and H3K36me3 at the *VRN1* locus during embryogenesis plays a critical role in maintaining the repression of *VRN1* prior to cold exposure, particularly the accumulation of H3K27me3 during embryogenesis.

### Discovery of novel regulators for vernalization response in winter wheat

Drawing insights from the transcriptional profiles of *VRN1* and *FLC* throughout the life cycle of wheat and *Arabidopsis* (27) and considering the reciprocal regulation by H3K27me3 and H3K36me3 in vernalization response, maintenance, and resetting (21, 22), we hypothesized that genes exhibiting analogous patterns of transcriptional and chromatin state changes might also play a role in vernalization-regulated flowering.

The transcriptional level of *VRN1* steadily increases during vernalization (from S0 to S4) and remains elevated post-vernalization (from S5 to S6) (Fig. 2E). Consequently, we conducted an exploration for *VRN1*-pattern-like genes using transcriptome, H3K27me3, and H3K36me3 data spanning S0 to S6. This effort resulted in the identification of 212 *VRN1*-pattern-like genes, discerned from the DEGs of C2 and C4 via K-means clustering (Table S4). Similar to *VRN1*, these genes exhibit consistent upregulation during vernalization and maintain heightened transcriptional activity after vernalization (Fig. 5A), all while being subject to regulation by both H3K27me3 and H3K36me3 (Fig. 5B). GO enrichment analysis reveals their involvement in flower development, signal transduction, auxin signaling, and shoot system development (Fig. 5C). Notably, the transcription factors within the *VRN1*-pattern-like genes are enriched in the MIKC_MADS family (Fig. 5D), particularly in the same subclade as *VRN1* (Fig. S5, Table S5). Among these genes, 93 harbor SNPs in their promoters or gene bodies significantly associated with heading time (Table S6), and 4 genes with severe TILLING mutations (Wang et al., 2023) (Table S7) exhibit aberrant flowering (Fig. 5E). These include the known flowering regulators TaFUL3 (43) and TaSOC1 (47), which display strikingly similar transcriptional and epigenetic profiles to *VRN1* (Fig. 5F). Notably, seven SNPs (red line in Fig. 5F) associated with the heading time are located in the promoter and gene body of *TaFUL3_2A* (Fig. 5G, Table S8).

**Fig. 5.**
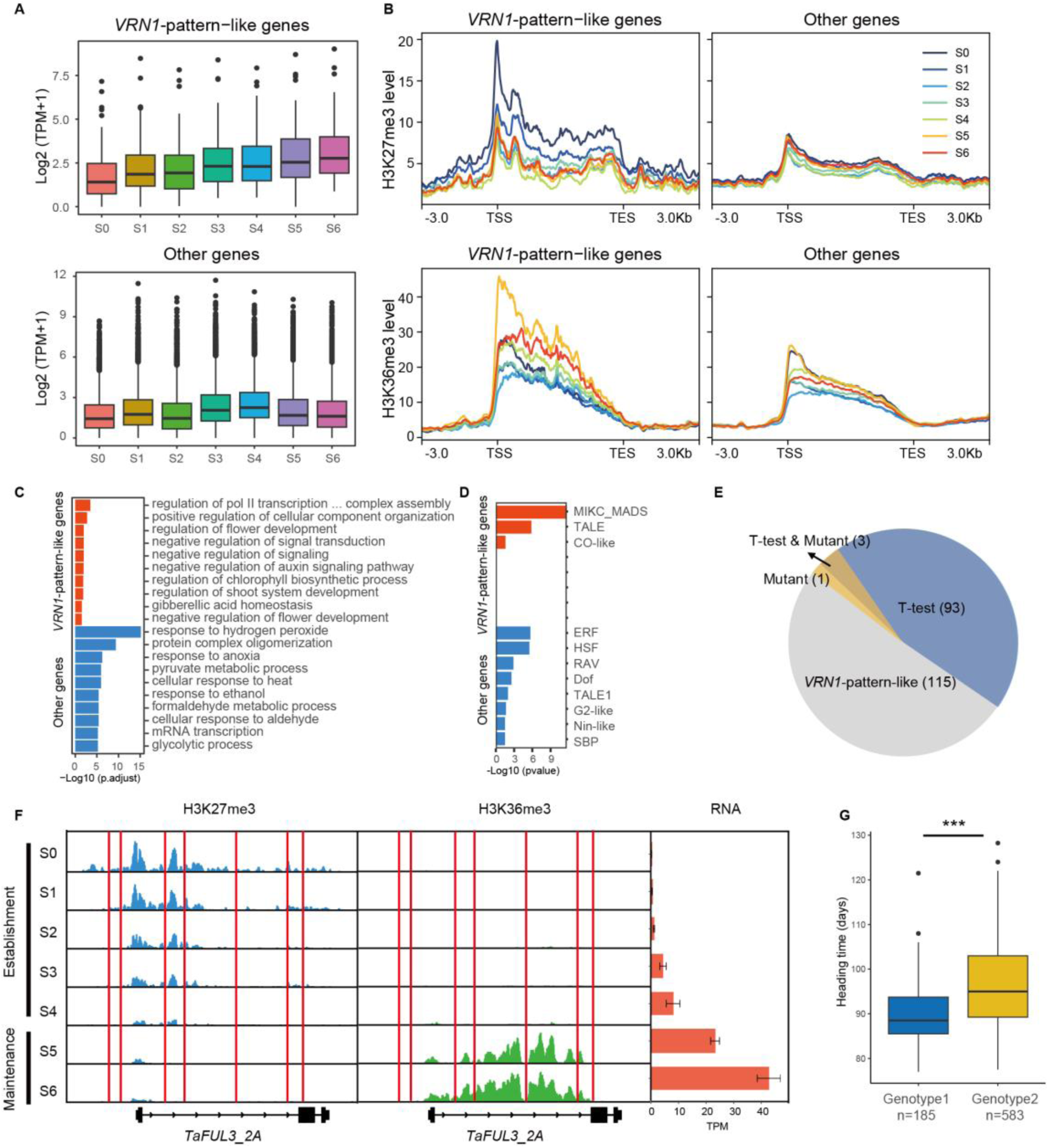
Identification and characterization of *VRN1*-pattern-like gene. **A.** The transcription pattern of *VRN1*-pattern-like genes in C2 and C4. Other refers to other genes in C2 and C4. **B.** The H3K27me3 and H3K36me3 dynamic of *VRN1*-pattern-like genes in C2 and C4. Other refers to other genes in C2 and C4. **C.** The GO enrichment analysis of *VRN1*-pattern-like genes and other genes in C2 and C4. **D.** The TF family enrichment analysis of *VRN1*-pattern-like genes and other genes in C2 and C4. **E.** Different types of *VRN1*-pattern-like genes. T-test represents SNP sites on the promoter or genebody of the gene that are significantly associated with heading time. Mutant represents an abnormal flowering phenotype in plants with this gene mutation. **F.** The H3K27me3, H3K36me3 and transcriptional level changes of *TaFUL3_2A* from S0 to S6. **G.** The heading time of two haplotypes divided by seven SNPs (red line in F) within promoter and genebody of *TaFUL3_2A*.

FLC, a flowering suppressor, plays a central role in mediating vernalization-regulated flowering in *Arabidopsis* (48). Consequently, efforts in previous research aimed to pinpoint its homologous gene in wheat (34, 49). We also found that these genes show high sequence similarity to *FLC* (highlighted in green) (Fig. S6), and the transcriptional inhibition of these genes by vernalization (Fig. S5, Table S5). However, the H3K27me3 and H3K36me3 level are not aligned with the transcription during vernalization (Fig. 6A).

**Fig. 6.**
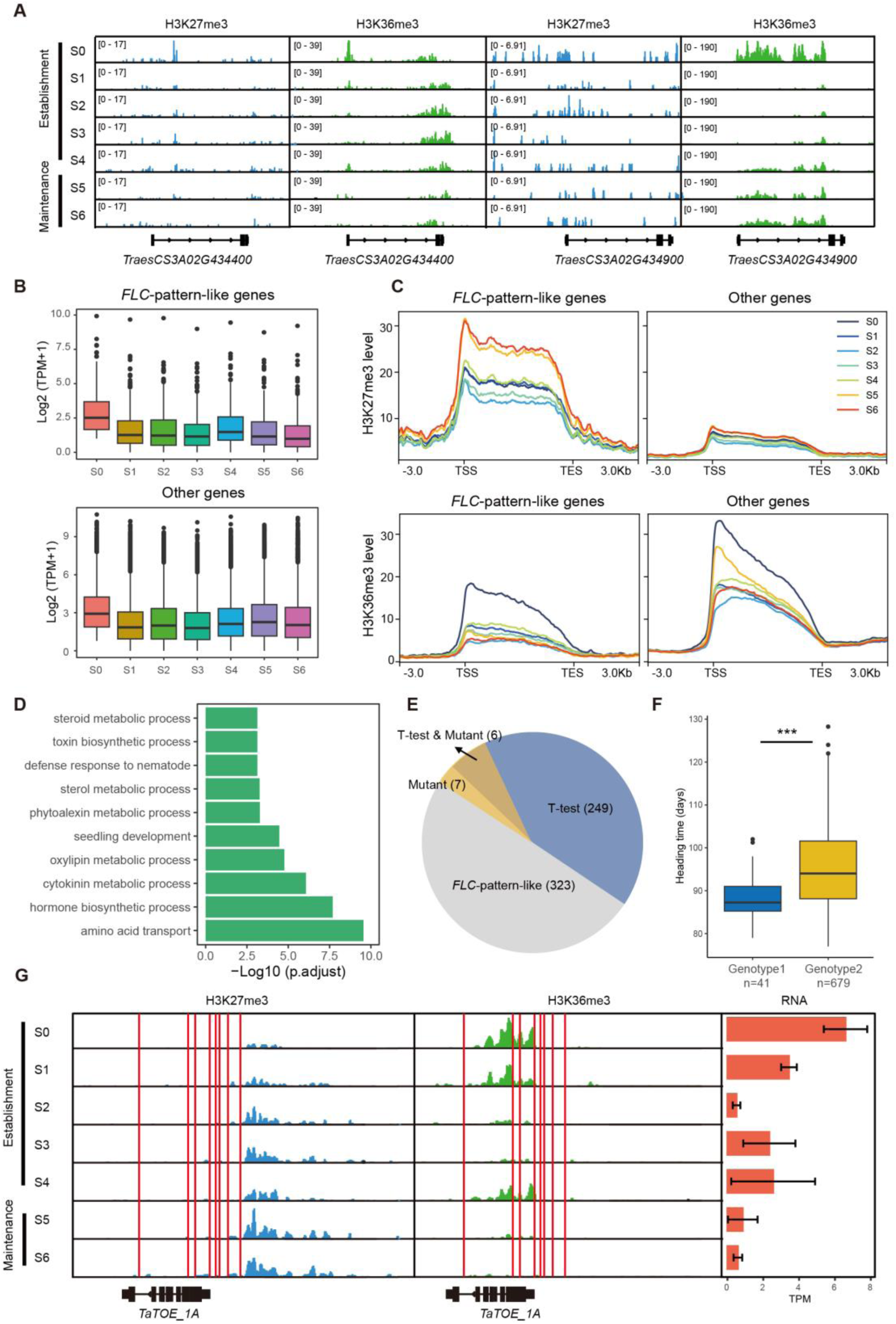
Identification and characterization of *FLC*-pattern-like gene. **A.** The H3K27me3 and H3K36me3 dynamic of FLC orthologs reported in previous study (85). **B.** The transcription pattern of *FLC*-pattern-like genes in C5. Other refers to other genes in C5. **C.** The H3K27me3 and H3K36me3 dynamic of *FLC*-pattern-like genes in C5. Other refers to other genes in C5. **D.** The GO enrichment analysis of *FLC*-pattern-like genes. **E.** Different types of *FLC*-pattern-like genes. T-test represents SNP sites on the promoter or genebody of the gene that are significantly associated with heading time. Mutant represents an abnormal flowering phenotype in plants with this gene mutation. **F.** The heading time of two haplotypes divided by eight SNPs (red line in G) within promoter and genebody of *TaTOE_1A*. **G.** The H3K27me3, H3K36me3 and transcriptional level changes of *TaTOE_1A* from S0 to S6.

Giving the importance of H3K27me3 and H3K36me3 in shaping *FLC* transcription pattern (21, 22), we embarked on a quest to identify *FLC*-pattern-like genes in other gene families using transcriptome, H3K27me3, and H3K36me3 data. Through this exploration, we discovered 585 *FLC*-pattern-like genes within DEGs of C5 through K-means clustering (Table S4). These genes undergo transcriptional inhibition by vernalization (Fig. 6B) and are associated with increased H3K27me3 and decreased H3K36me3 (Fig. 6C). They are linked to various biological processes, including steroid metabolic processes, toxin biosynthesis, seedling development, and hormone biosynthesis (Fig. 6D). Within this set, 249 genes (Table S6) harbor SNPs significantly associated with flowering time, while 13 genes (Table S7) with severe TILLING mutations can exert an impact on wheat flowering time (Fig. 6E). Furthermore, *TaTOE1* has been previously reported to influence wheat flowering (50) and we identified eight SNPs associated with heading time within its promoter and genebody (Fig. 6F, G, Table S9).

In sum, through a thorough examination of transcriptional and epigenetic feature similarities, combined with available genotype and phenotype at population level and TILLING mutant information, we have identified potential novel regulators governing vernalization-regulated flowering in winter wheat.

## Discussion

Plants respond to seasonal temperature changes, notably extended winter cold, orchestrating crucial developmental shifts such as flowering (1). Unlike in *Arabidopsis*, where cold silences the flowering repressor *FLC*, cereals like wheat activate the flowering promoter *VRN1* during winter (3, 51). Despite their opposite roles in flowering regulation, our in-depth analysis of transcriptome and epigenetic profiles revealed a conserved epigenetic regulatory model for mediating *VRN1* transcriptional dynamic in winter wheat, akin to *FLC* regulation in *Arabidopsis*.

*FLC*, active pre-cold, is gradually silenced, kept low post-warm return (52, 53). It remains low throughout gametophyte generation, reactivates during embryogenesis, and stays high during seedling development, resetting the vernalization requirement for the next generation (25–27). In our study, transcriptome analysis of the shoot apex during the life cycle of winter wheat KN9204 indicated that *VRN1* is initially inactive in seedlings before cold treatment. During the cold winter, *VRN1* gradually activates and remains high in the following spring in the inflorescence tissue (Fig. 2). Subsequently, *VRN1* is re-silenced during embryogenesis to reset the vernalization requirement (Fig. 4).

In *Arabidopsis*, *FLC*’s chromatin status correlates with its transcription. Pre-cold, H3K4me3 and H3K36me3 mark *FLC*, shifting to H3K27me3 at the nucleation site during cold, spreading across the entire gene body post-warm return, serving as a memory of the vernalization treatment. Similarly, we observed a conserved yet mirrored pattern of H3K27me3 and H3K36me3 dynamics at the *VRN1* locus in winter wheat (Fig. 2). H3K27me3 decreases, H3K36me3 but not H3K4me3 accumulates during cold, persisting post-warm return (Fig. 2). *T*a*sdg8-cr* lines fail to sustain active *VRN1* expression post-vernalization, resulting in delayed flowering (Fig. 3). During embryogenesis, H3K36me3 reduction correlates with *VRN1* transcription drop in both winter and spring wheat, while H3K27me3 accumulates selectively in winter wheat, crucial for inhibiting unlicensed *VRN1* transcription pre-cold in *Tafie-cr-87* (Fig. 4). It would be intriguing to unravel the mechanisms underlying the dynamic patterns of H3K27me3 and H3K36me3 at the *VRN1* locus throughout the life cycle of winter wheat. Furthermore, exploring the association between the dynamics of the epigenetic landscape and transcriptional patterns across different winter and spring *VRN1* alleles presents a compelling avenue for future analysis.

Giving *FLC*’s crucial role in vernalization responses across species, substantial efforts have been made to find *FLC* counterparts in monocots (34, 54). Based on sequence similarity, TaOS2/TaAGL33 has been identified as a potential floral repressor in wheat, exhibiting a response to cold exposure, albeit with a less pronounced early flowering phenotype (33, 55). Similarly, in barley and *Brachypodium*, BdOS2 and HvOS2 also function as floral repressors (33, 56). Our analysis, however, didn’t reveal a correlated H3K27me3 and H3K36me3 pattern at the TaOS2 locus during development (Fig. 6), suggesting it might not be an epigenetic regulation target for vernalization. Recognizing the mitotic memory in vernalization treatment, we suggest that epigenetic modification patterns could be vital markers for identifying genes similar to *FLC* and *VRN1*. By integrating expression patterns and H3K27me3/H3K36me3 profiles during vernalization (Fig. 5,6), we identified potential factors like TaFUL3 and TaTOE1, associated with flowering and showing natural variations linked to heading time diversity in extensive wheat populations, meriting further analysis.

As global warming alters temperature conditions (57), impacting the growth areas for wheat varieties, it becomes crucial to understand and adapt vernalization responses to ensure optimal grain yield. Fine-tuning the major vernalization response locus *VRN1*, along with the identification of additional genetic loci controlling the vernalization response, could enhance wheat’s resilience to environmental change.

## Materials and Methods

### Growth conditions, vernalization treatment and experimental materials

The winter wheat cultivar KN9204 used in this study were grown during the normal growing seasons at the Experimental Station of Institute of Genetics and Developmental Biology, Chinese Academy of Sciences, Beijing, China at the beginning of October each year. They experienced natural vernalization during the winter and flowered the following spring.

The stage-specific apexes of wheat (S0-S6) were dissected under the stereomicroscope based on the anatomic and morphological features at the right time (Fig. 1A), and immediately frozen in liquid nitrogen and stored at -80 °C. About 10 to 50 spikes were pooled. The stamens were removed before the pollen maturation. Then, we conducted artificial pollination and recorded the number of days to ensure the accurate time of seed development. The seeds at DAP0, DAP2, DAP6, DAP8, and DAP16 stage were sampled for later use of total RNA extraction and nuclei isolation. The dissection method of embryo and embryo sac follow the instructions in the previous article (37). Embryo sacs containing egg/embryo and surrounding tissue in DAP0 and DAP2 were dissected in a 5% Sucrose solution containing 0.1% RNase inhibitor with fine forceps using the dissecting microscope. For DAP6 and later stages, the independent embryo was detached from the embryo sac using the same method. Embryos and embryo sacs sampled from eight to ten spikes were pooled for one biological replicate in early stages and three to five were pooled for one biological replicate in late stages. The RNA-seq (three replicates), ATAC-seq and CUT&Tag (two replicates) experiments at twelve development stages were carried out.

For artificial vernalization treatment, seeds of winter wheat and spring wheat (*Triticum aestivum* cv. KN9204, KN199 and CS), seeds of *TaFIE* or *TaSDG8* transgenic lines were surface sterilized in 2% NaClO for 20 minutes, then rinsed overnight with flowing water. The seeds were germinated and then transferred to soil, and grown at 4°C in the dark for 0, 20, 30 days (V0, V20, V30) and then grown at room temperature for 10 days (V30+10). Wheat leaves from these samples were used to extract RNA for RT–qPCR or isolate nuclei for CUT&Tag–qPCR. Then were transferred in a greenhouse (20°C–22°C, 16-h light/8-h dark). Finally, the heading time is counted, which is the time from sowing to heading minus the corresponding 4°C vernalization treatment time.

### TILLING mutation library and identification of lines with heading time difference

The previously published KN9204 TILLING mutation library (58) was used to screen mutants with differences in heading time from wild-type KN9204. Each mutant line was planted in a 1.5 m single-row plot with 10cm between plants and 25 cm between rows and managed according to local practices. Heading time was calculated as the days from sowing to half of the spikes in a row heading.

### RNA-seq, ATAC-seq and CUT&Tag experiment

Total RNA was extracted using HiPure Plant RNA Mini Kit (Magen, R4111-02). RNA-seq libraries’ construction and sequencing platform were the same as previous description (37), by Annoroad Gene Technology.

ATAC-seq and CUT&Tag experiments followed the previously described method (37). Tn5 transposase used and tagmentation assay is done following the manual (Vazyme, TD501-01). Libraries were purified with AMPure beads (Beckman, A63881) and sequenced using the Illumina Novaseq platform at Annoroad Gene Technology. Antibodies used for histone modifications are listed in Table S10.

### RNA-seq data analysis

Fastp (v0.23.1) (59) was used to process raw reads to remove adapters, low quality bases and reads. Then the cleaned reads were aligned to IWGSC RefSeq v1.0 (55) using hisat2 (v2.0.5) (60). Samtools (v1.5) (61) was used to sort the aligned reads and converted it into bam files. The gene count matrix was calculated by featureCounts (v2.0.1) (62) with parameter “-p –P –B -C” The counts matrixs were used as inputs to identify differentially expressed genes using R package DESeq2 (v1.28.1) (63), with a threshold of p.adj < 0.05 and abs(log2FoldChange) > 1. TPM (Transcripts Per Kilobase Million) values generated from the counts matrix were used to characterize gene expression and used for hierarchical clustering analysis. K-means cluster analysis of differentially expressed genes were performed in R (v4.0.2) using Z-scaled TPM and the gene expression heatmap was displayed using R package ComplexHeatmap (v2.4.3) (64). The gene ontology enrichment analysis was done on using R package clusterProlifer (v3.16.1) (65) and the gene ontology annotation was performed using PANTHER (66). The identification and family classification of wheat TFs were predicted on the PlantTFdb website (http://planttfdb.gao-lab.org/) (67). Enrichment analysis of TF families for VRN1-like genes was achieved using the enricher function in R package clusterProlifer (v3.16.1) (65).

### ATAC-seq and CUT&Tag data analysis

Raw reads were processed by fastq (v0.23.1) with “--detect_adapter_for_pe” parameter to remove adapters, trim low quality bases and filter bad reads. Clean reads were aligned to IWGSC RefSeq v1.0 using bwa mem algorithm (v 0.7.17) (68) with “-M” parameter. The aligned reads were sorted and filtered by samtools (v1.5) with “-F 1804 -f 2 -q 30” parameter. Picard (v 2.23.3) was used to remove the duplicated reads. The de-duplicated bam files from two biological replicates were merged by samtools (v1.5) and converted into bigwig files using bamCoverage provided by deeptools (v3.5.0) (69) with parameters “-bs 10 --effectiveGenomeSize 14600000000 --normalizeUsing RPKM --smoothLength 50”. The bigwig files were visualized using deeptools (v3.5.0) and IGV (v2.8.0.01) (70).

The peak calling was done using merged bam files by macs2 (v 2.2.7.1) (71). For ATAC-seq data, the parameter of peak calling using macs2 is “-q 0.05 -f BAMPE --nomodel -- extsize 200 --shift -100 -g 14600000000”. For narrow peak of H3K27ac, H3K4me3 and H2A.Z, the peak calling parameter is “-p 1e-3 -f BAMPE -g 14600000000 --keep-dup all”. For broad peak of H3K27me3 and H3K36me3, the peak calling parameter is “-f BAMPE - g 14600000000 --keep-dup all --broad --broad-cutoff 0.05”. Peaks from S0 to S6 of the same data type were merged using bedtools (v2.29.2) (72) to generate the reference peaks. Peak was annotated to the wheat genome using the R package ChIPseeker (v1.24.0) (73).

For quantification of ATAC-seq and CUT&Tag data, reads count under reference peak and the normalized CPM (DBA_SCORE_TMM_READS_EFFECTIVE_CPM) values were generated using R package DiffBind (v2.16.2). The CPM values of reference peak were used to do hierarchical clustering analysis.

### Chromatin state analysis

ChromHMM (v1.22) (40) was used to train models containing 5 to 25 chromatin states using all CUT&Tag data. CompareModels was used to calculate correlation between different models. OverlapEnrichment was used to calculate the enrichment of each state at different locations in genome.

### Association analysis between genotype and heading time

The genotype and heading time data from 788 wheat cultivars was obtained from the published data (74). We extracted SNPs located in the promoter and genebody of VRN1-pattern-like and FLC-pattern-like genes. The population was divided into two sub-population by each single SNP, and t-test was performed to test the significance of the difference in heading time between two sub-population. The significance threshold is set to 1e-6.

### Phylogenetic tree of MADS family

All the proteins in *Arabidopsis thaliana*, wheat, rice, barley, and *Brachypodium distachyon* that were predicted by hmmsearch (v3.3.2) (75) to contain the MADS domain were considered to be the proteins of the MADS family. The longest protein of the MADS family gene was aligned by muscle (v3.8.31) (76), and then trimal (v1.4.rev15) (77) was used to cut the aligned sequences. Only the sequence that existed in 80% of the genes was retained. RAxML (v8.2.12) (78) was used to establish the phylogenetic tree of the MADS family.

### Construction and genotyping of CRISPR/Cas9 lines

The construction and identification of *Tafie-cr-87* mutants was reported in previous study (37). For mutant *Tasdf8-cr-3/5*, plasmid constructs pJIT163-Ubi-Cas9 and Pu6-gRNA39 were used (79, 80). Gene-specific primers were designed around the gRNA target site to identify the *Tasdg8-cr-3/5* mutants. PCR products were genotyped by Sanger sequencing. Primers used for construction and identification are listed in Table S11.

### In situ hybrization

RNA in situ hybridization was carried out as described previously (81). Fresh young spikes and seeds were fixed in formalin-acetic acid-alcohol overnight at 4 °C, dehydrated through a standard ethanol series, embedded in Paraplast Plus tissue-embedding medium (Sigma-Aldrich, P3683), and sectioned at 8-10 μm width using a microtome (Leica Microsystems, RM2235). Digoxigenin-labeled RNA probes based on the sequence of *VRN1* were synthesized using a DIG northern Starter Kit (Roche, 11277073910), according to the manufacturer’s instructions. Primers used for probe synthesis are listed in Table S11.

### RT-qPCR

First-strand cDNA was synthesized from 2 μg of DNase I-treated total RNA using the TransScript First Strand cDNA Synthesis SuperMix Kit (TransGen, AT301-02). RT-qPCR was performed using the ChamQ Universal SYBR qPCR Master Mix (Vazyme, Q711-02) by QuantStudio5 (Applied biosystems). The expression of interested genes was normalized to Tubulin for calibration, and the relative expression level is calculated via the 2-ΔΔCT analysis method (82). Primers used for RT-qPCR are listed in Table S11.

### CUT&Tag-qPCR

CUT&Tag-qPCR was performed as previously reported with minor modifications (83). Briefly, the DNA products of CUT&Tag were divided, 6µL was used as “Input”, and the other 18µL undergoing library PCR amplification and purification was used as “IP products.” Finally, the “Input” and “IP products” were dilute 30 times for qPCR assay. The primers for specific regions are provided in Table S11.

## Data availability

The omics data at S0, S2, S3, DAP0, DAP2, DAP6 stages are reported for the first time in this study and the raw sequence data have been deposited in the Genome Sequence Archive of the National Genomics Data Center, China National Center of Bioinformation/Beijing Institute of Genomics, Chinese Academy of Sciences (<u>PRJCA022101</u>). Other omics data were download from our previous report (84).

## Acknowledgments

This research was supported by National Natural Science Foundation (31970529), Beijing Natural Science Foundation Outstanding Youth Project (JQ23026), National Key Research and Development Program of China (2021YFD1201500), the Major Basic Research Program of Shandong Natural Science Foundation (ZR2019ZD15).

## Figures

**Fig. S1.**
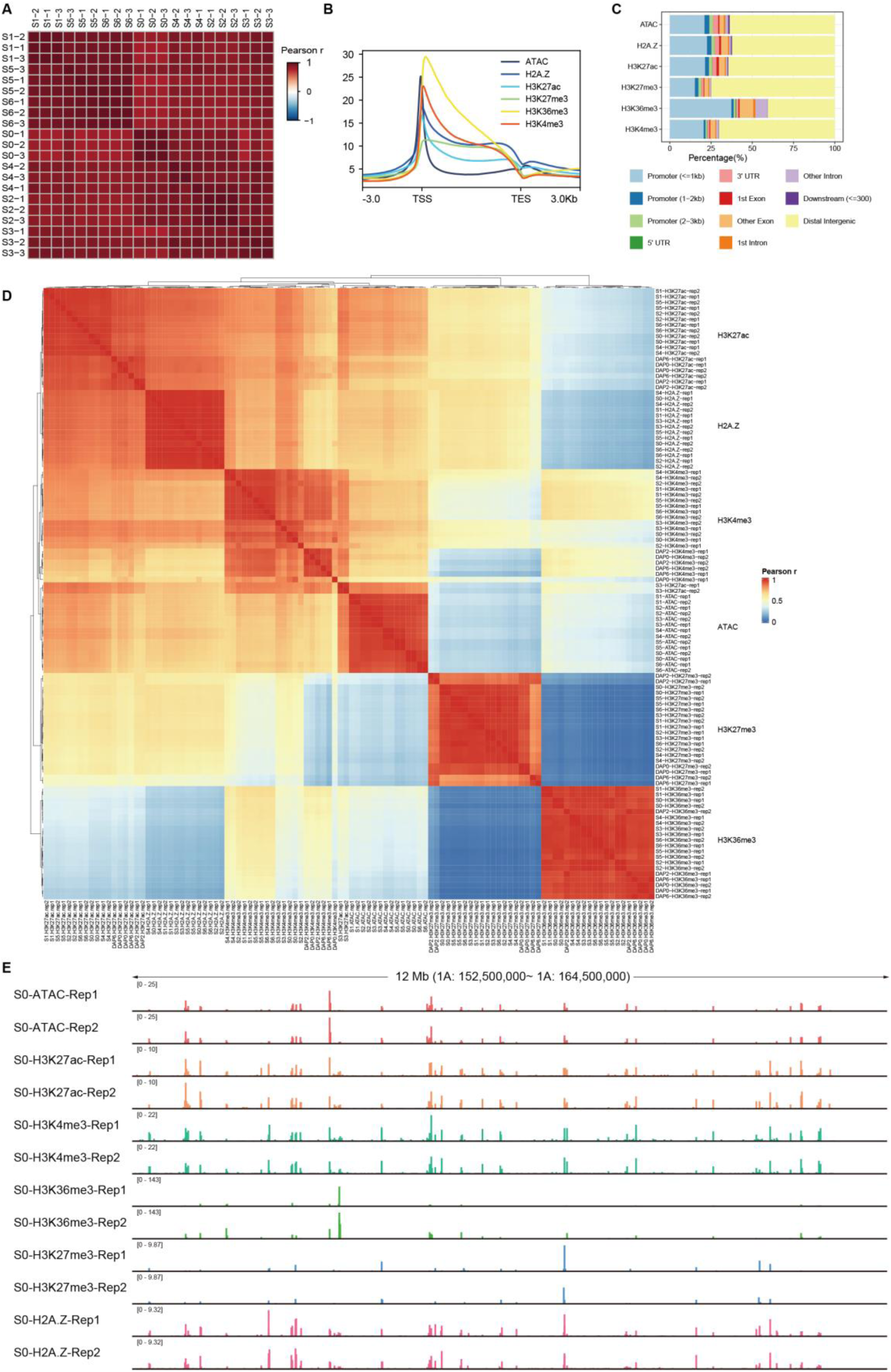
Quality control for transcriptome and epigenome data. **A.** Pearson correlation analysis of all transcriptome data. The color represents Pearson correlation coefficient. **B.** Epigenetic profile on genes, taking S0 as an example. TSS: transcription start site; TES: transcription end site. **C.** The distribution of epi-genome peaks in the genome, taking S0 as an example. **D.** Pearson correlation analysis of all epi-genome data. The color represents Pearson correlation coefficients.

**Fig. S2.**
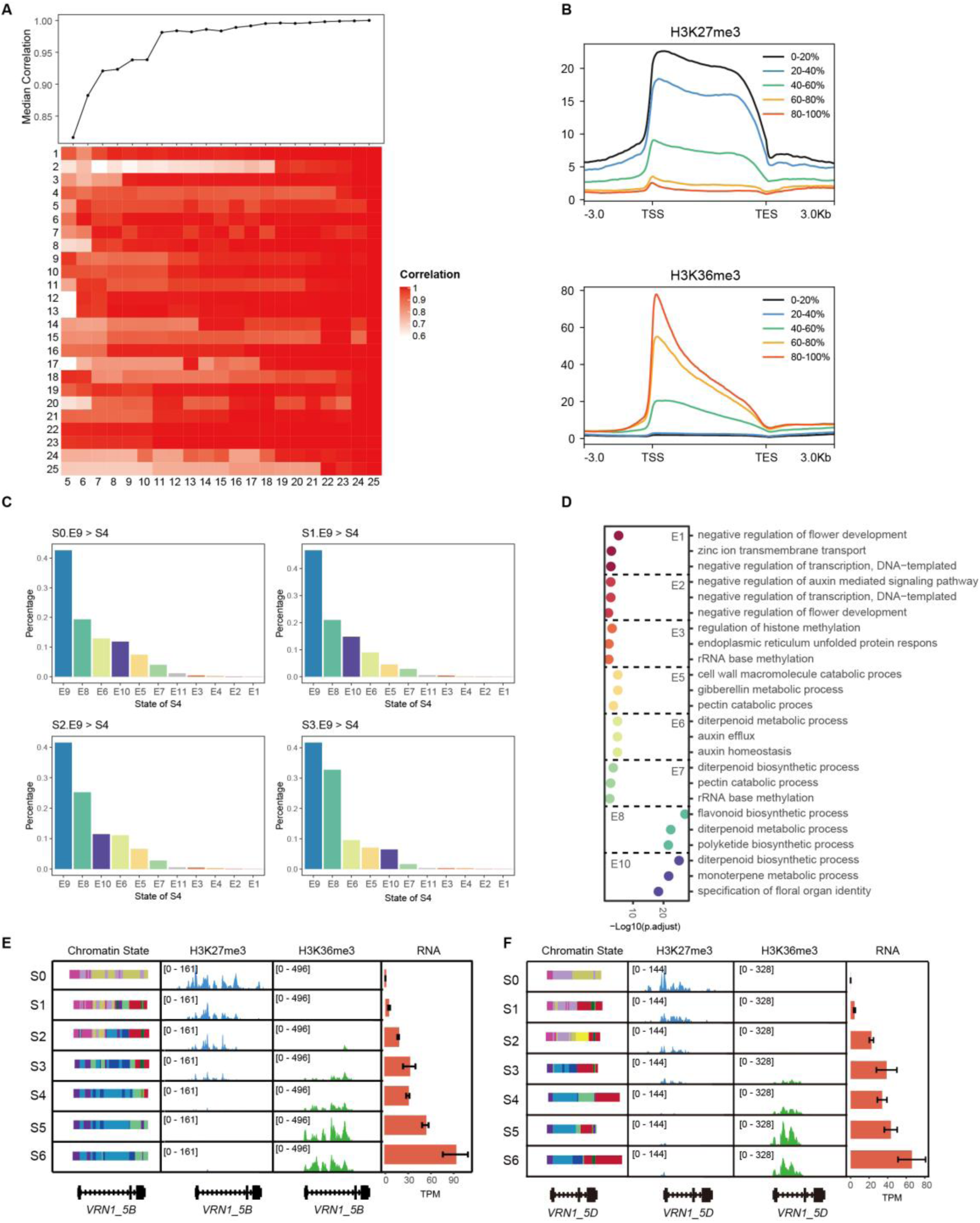
Changes of individual chromatin state during vernalization. **A.** Correlation of different chromatin state models trained by chromHMM. The color represents the level of correlation. **B.** The levels of H3K27me3 (top) and H3K36me3 (bottom) in genes with different transcription levels, take S0 as an example. Different colors represent genes at different levels of transcription. All genes in the S0 period were ranked from lowest to highest expression, with 0-20% representing the top 20% with the lowest expression and 80-100% representing the top 20% with the highest expression. **C.** Changes in E9 state from S0-S3 to S4 stage. **D.** The GO enrichment analysis of genes with altered E9 states from S0-S3 to S4 stage. **E, F.** The chromatin state, H3K27me3, H3K36me3 and transcriptional level changes of *VRN1_5B* (E) and *VRN1_5D* (F) from S0 to S6.

**Fig. S3.**
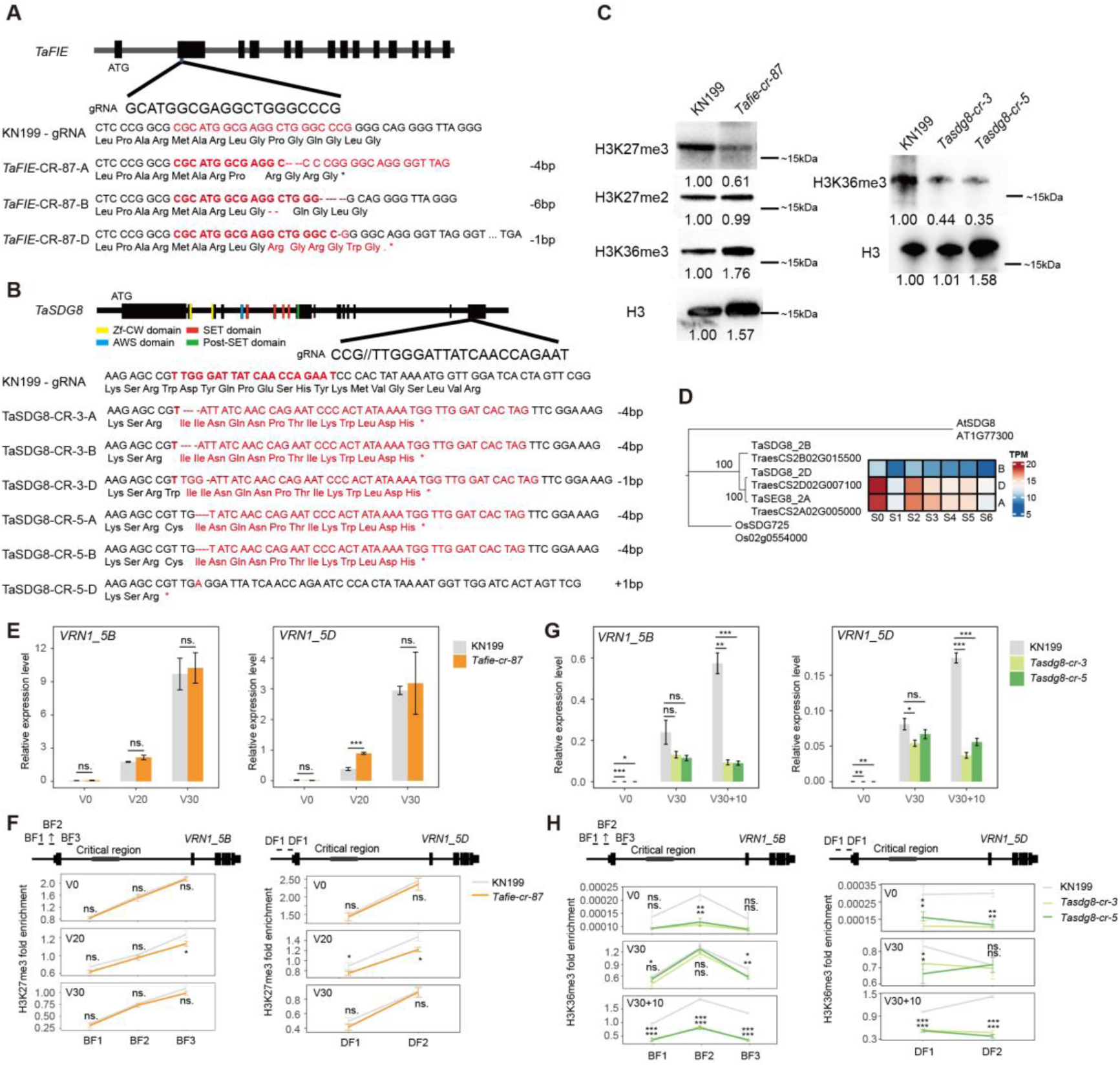
Generation of *Tafie* and *Tasdg8* crispr/cas9 lines. **A.** Indication of genome editing sites and genotyping of *Tafie* crispr/cas9 line. **B.** Indication of genome editing sites and genotyping of *Tasdg8* crispr/cas9 lines. **C.** The western blot shows the H3K27me3, H3K27me2, and H3K36me3 level in wild type (KN199) and *Tafie, Tasdg8* crispr/cas9 lines (*Tafie-cr-87 and Tasdg8-cr-3/5*). The number show quantification of protein levels compared to the wild type (KN199) in each condition. **D.** The phylogenetic tree and expression level of *TaSDG8*. **E.** The relative expression level of *VRN1_5B* and *VRN1_5D* in seedlings of wild type (KN199) and *Tafie* crispr/cas9 (*Tafie-cr-87*) line under different vernalization treatments. **F.** The H3K27me3 level at different location of *VRN1_5B* and *VRN1_5D* in seedlings of wild type (KN199) and *Tafie* crispr/cas9 line (*Tafie-cr-87*) at heading stage under different vernalization treatments. **G.** The relative expression level of *VRN1_5A* and *VRN1_5D* in seedlings of wild type (KN199) and *Tasdg8* crispr/cas9 (*Tasdg8-cr-3* and *Tasdg8-cr-5*) lines under different vernalization treatments. **H.** The H3K36me3 level at different location of *VRN1_5B* and *VRN1_5D* in seedlings of wild type (KN199) and *Tasdg8* crispr/cas9 (*Tasdg8-cr-3*, *Tasdg8-cr-5*) lines under different vernalization treatments.

**Fig. S4.**
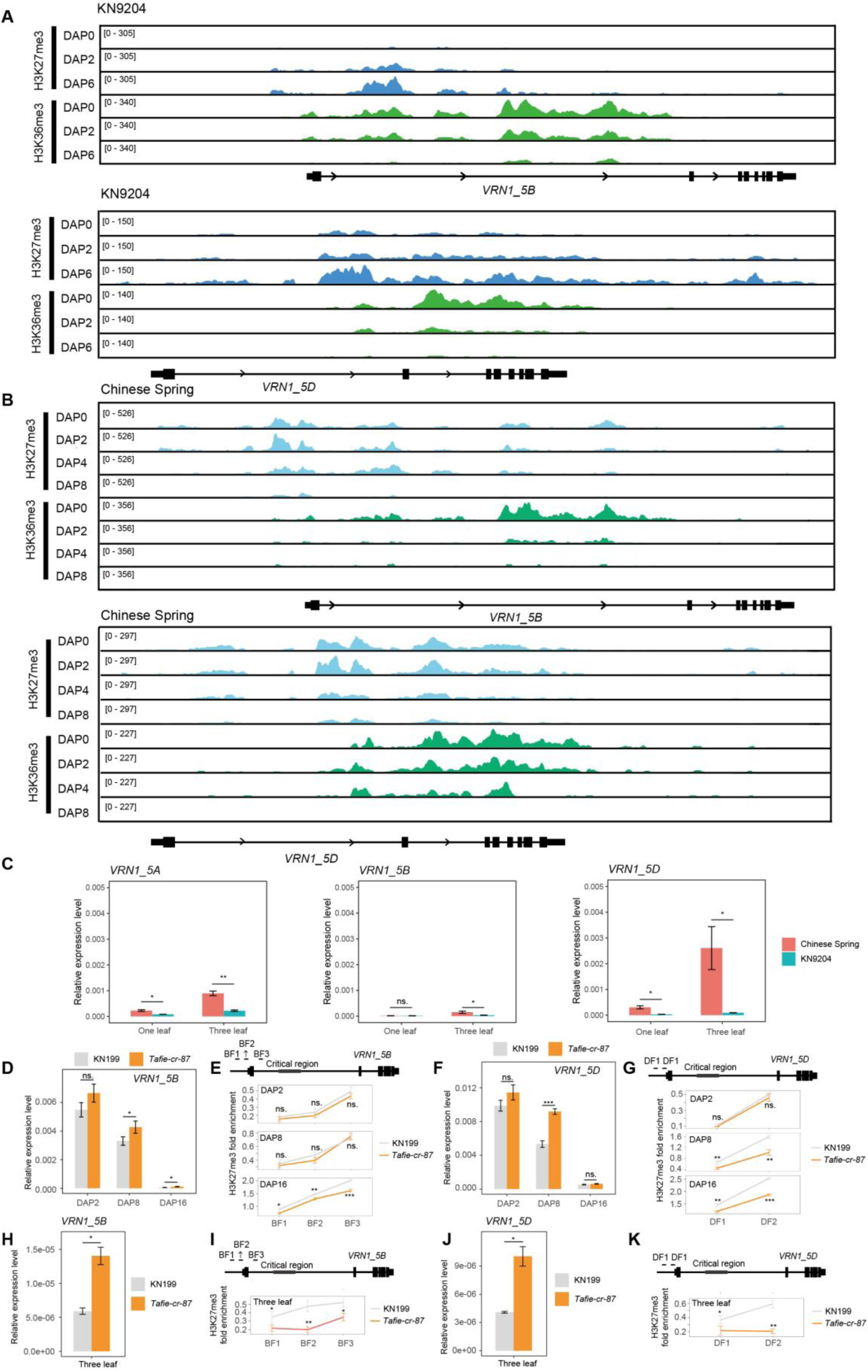
mRNA level and H3K27me3, H3K36me3 abundance of VRN1_5B and VRN1_5D during embryogenesis. **A.** The H3K27me3 and H3K36me3 changes of *VRN1_5B* and *VRN1_5D* during embryogenesis in winter wheat KN9204. **B.** The H3K27me3 and H3K36me3 changes of recessive allele *VRN1_5B* and dominant allele *VRN1_5D* during embryogenesis in spring wheat CS. **C.** The relative expression level of *VRN1* in leaves of CS and KN9204. **D, H**. The relative expression level of *VRN1_5B* in embryos and leaves of wild type (KN199) and *Tafie* crispr/cas9 (*Tafie-cr-87*) line. **E, I.** The H3K27me3 level at different location of *VRN1_5B* in embryos and leaves of wild type (KN199) and *Tafie* crispr/cas9 line (*Tafie-cr-87*). **F, J.** The relative expression level of *VRN1_5D* in embryos and leaves of wild type (KN199) and *Tafie* crispr/cas9 (*Tafie-cr-87*) line. **G, K.** The H3K27me3 level at different location of *VRN1_5D* in embryos and leaves of wild type (KN199) and *Tafie* crispr/cas9 line (*Tafie-cr-87*).

**Fig. S5.**
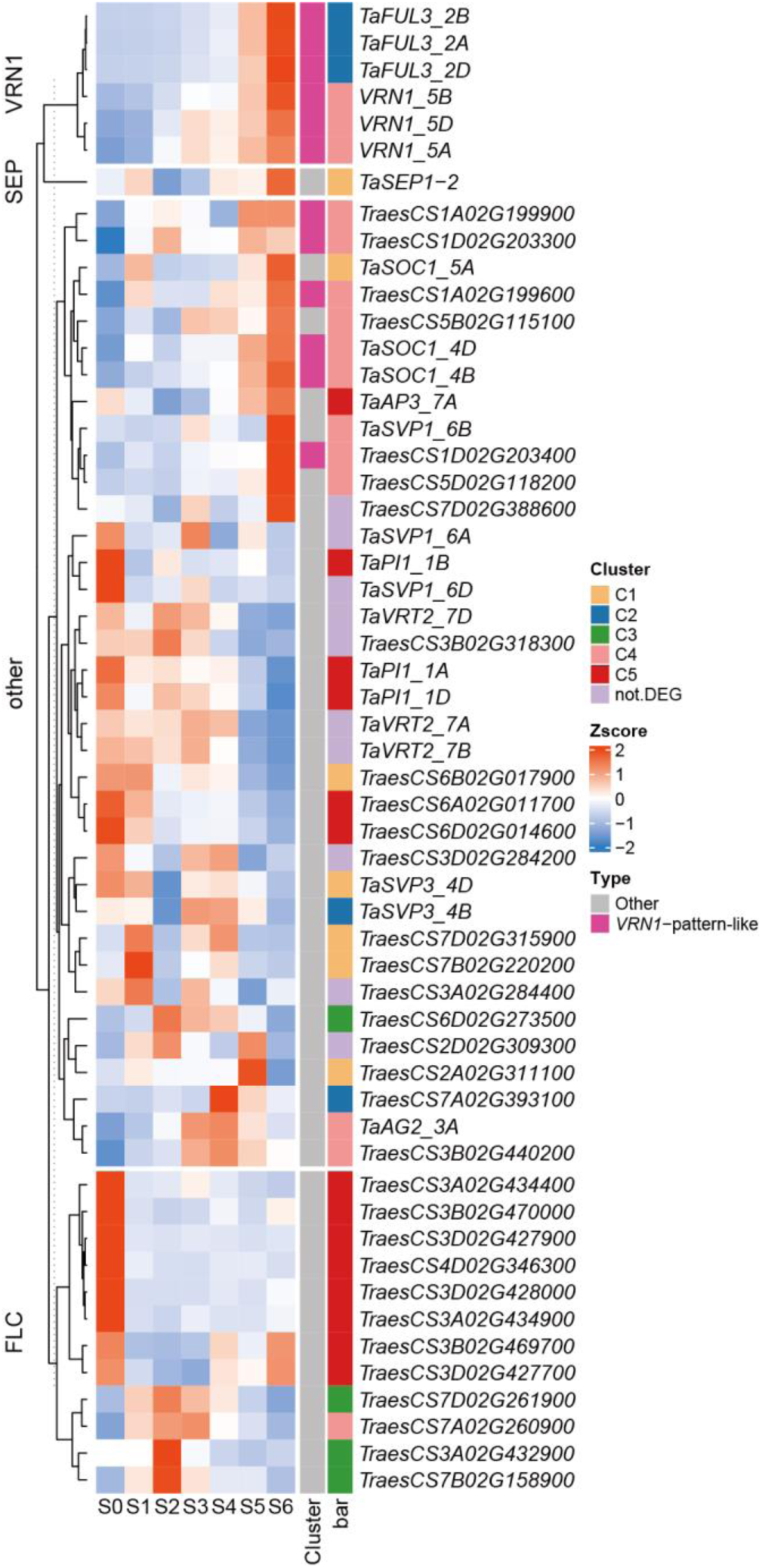
Expression pattern of MADS genes from S0 to S6. VRN1 represents genes in *VRN1* subfamily, SEP represents genes in *SEP* subfamily, FLC represents genes in *FLC* subfamily and other represents genes in other subfamily of MADS.

**Fig. S6.**
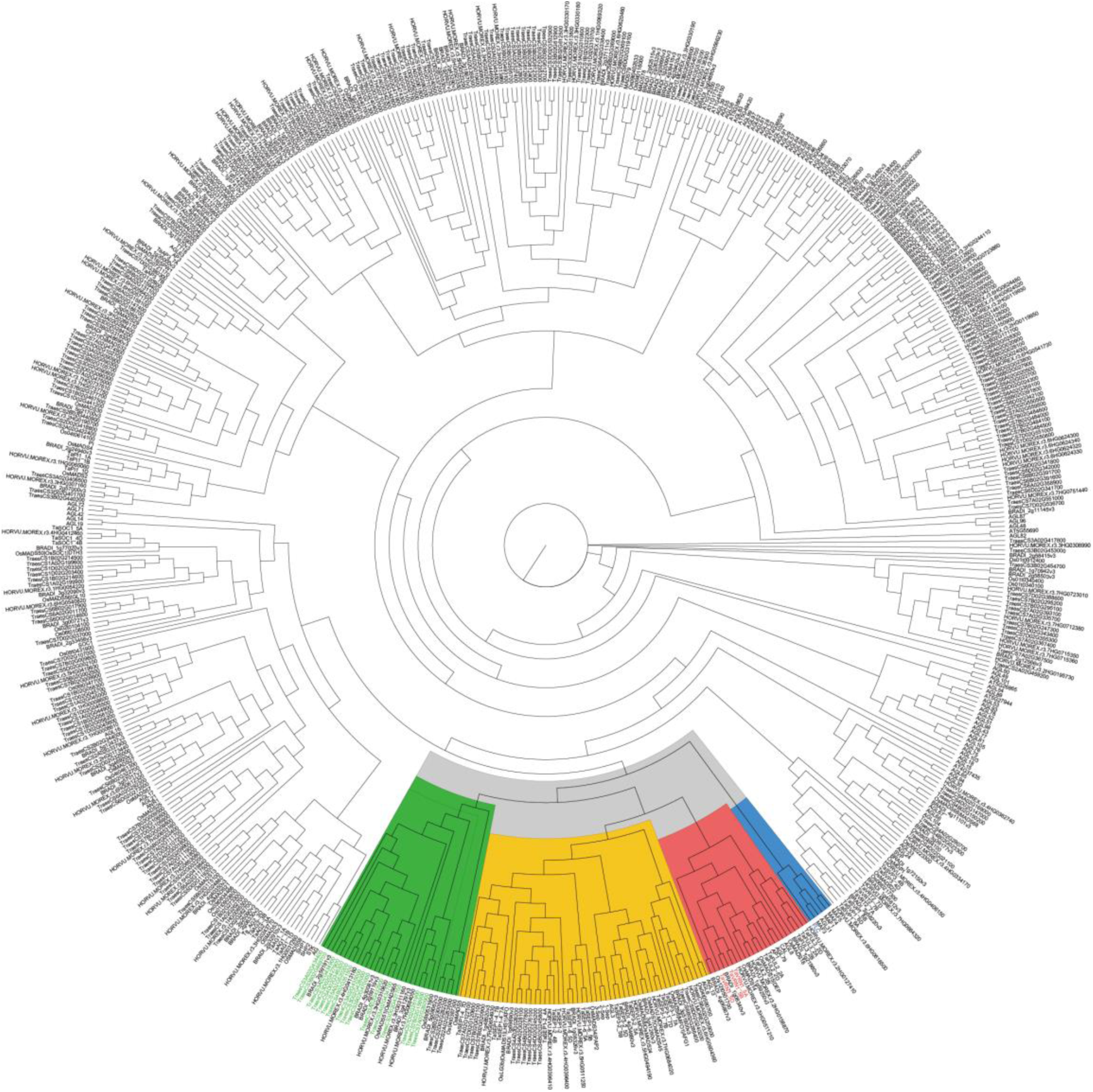
Phylogenetic tree of the MADS family from *Arabidopsis thaliana*, wheat, rice, barley and *Brachypodium distachyon*. Blue represents the subfamily of Arabidopsis *FLC*, red represents the subfamily of wheat *VRN1*, yellow represents the subfamily of *SEP,* and green represents the subfamily of the previously reported orthologs of *FLC* in wheat (85).

## Supplementary tables

**Table S1** Differentially expressed genes from S0 to S4.

**Table S2** The expression level of *ICE-CBF-COR* cluster genes.

**Table S3** The changes of state E9 from S0-S3 to S4.

**Table S4** The VRN1-pattern-like and FLC-pattern-like genes.

**Table S5** The expression level of genes in MADS family.

**Table S6** SNPs located within the promoter of genebody of *VRN1*-pattern-like or *FLC*-pattern-like genes that were significantly associated with heading time in GWAS.

**Table S7** Mutant lines containing severe mutations in *VRN1*-pattern-like or *FLC*-pattern-like genes.

**Table S8** SNPs associated with heading time located on the promoter and genebody of *TaFUL3*_2A.

**Table S9** SNPs associated with heading time located on the promoter and genebody of *TaTOE*_1A.

**Table S10** Antibodies used in this study.

**Table S11** Primers used in this study.

## References

1. P. Chouard, Vernalization and its Relations to Dormancy. Annual Review of Plant Physiology 11, 191–238 (1960).

2. R. Amasino, Seasonal and developmental timing of flowering. Plant J 61, 1001–1013 (2010).

3. S. Xu, K. Chong, Remembering winter through vernalisation. Nature Plants 4, 997–1009 (2018).

4. A. Distelfeld, C. Li, J. Dubcovsky, Regulation of flowering in temperate cereals. Current Opinion in Plant Biology 12, 178–184 (2009).

5. I. R. Henderson, C. Shindo, C. Dean, The Need for Winter in the Switch to Flowering. Annual Review of Genetics 37, 371–392 (2003).

6. L. Yan, et al., Positional cloning of the wheat vernalization gene VRN1. Proceedings of the National Academy of Sciences 100, 6263–6268 (2003).

7. L. Yan, et al., The wheat VRN2 gene is a flowering repressor down-regulated by vernalization. Science 303, 1640–1644 (2004).

8. L. Yan, et al., The wheat and barley vernalization gene VRN3 is an orthologue of FT. Proc Natl Acad Sci U S A 103, 19581–19586 (2006).

9. A. Chen, J. Dubcovsky, Wheat TILLING Mutants Show That the Vernalization Gene VRN1 Down-Regulates the Flowering Repressor VRN2 in Leaves but Is Not Essential for Flowering. PLOS Genetics 8, e1003134 (2012).

10. J. Xiao, et al., Wheat genomic study for genetic improvement of traits in China. Sci. China Life Sci. 65, 1718–1775 (2022).

11. D. Fu, et al., Large deletions within the first intron in VRN-1 are associated with spring growth habit in barley and wheat. Mol Genet Genomics 273, 54–65 (2005).

12. I. Diaz, et al., The GAMYB protein from barley interacts with the DOF transcription factor BPBF and activates endosperm-specific genes during seed development. Plant J 29, 453–464 (2002).

13. N. Kippes, et al., Identification of the VERNALIZATION 4 gene reveals the origin of spring growth habit in ancient wheats from South Asia. Proc Natl Acad Sci U S A 112, E5401–5410 (2015).

14. A. B. Shcherban, E. A. Salina, Evolution of VRN-1 homoeologous loci in allopolyploids of Triticum and their diploid precursors. BMC Plant Biol 17, 188 (2017).

15. Y. Chen, B. F. Carver, S. Wang, F. Zhang, L. Yan, Genetic loci associated with stem elongation and winter dormancy release in wheat. Theor Appl Genet 118, 881–889 (2009).

16. G. Li, et al., Vernalization requirement duration in winter wheat is controlled by TaVRN-A1 at the protein level. Plant J 76, 742–753 (2013).

17. J. Xiao, et al., O-GlcNAc-mediated interaction between VER2 and TaGRP2 elicits TaVRN1 mRNA accumulation during vernalization in winter wheat. Nat Commun 5, 4572 (2014).

18. S. N. Oliver, E. J. Finnegan, E. S. Dennis, W. J. Peacock, B. Trevaskis, Vernalization-induced flowering in cereals is associated with changes in histone methylation at the VERNALIZATION1 gene. Proc Natl Acad Sci U S A 106, 8386–8391 (2009).

19. S. N. Oliver, W. Deng, M. C. Casao, B. Trevaskis, Low temperatures induce rapid changes in chromatin state and transcript levels of the cereal VERNALIZATION1 gene. J Exp Bot 64, 2413–2422 (2013).

20. S. Xu, et al., The vernalization-induced long non-coding RNA VAS functions with the transcription factor TaRF2b to promote TaVRN1 expression for flowering in hexaploid wheat. Mol Plant 14, 1525–1538 (2021).

21. H. Yang, et al., Distinct phases of Polycomb silencing to hold epigenetic memory of cold in Arabidopsis. Science 357, 1142–1145 (2017).

22. H. Yang, M. Howard, C. Dean, Antagonistic roles for H3K36me3 and H3K27me3 in the cold-induced epigenetic switch at Arabidopsis FLC. Curr Biol 24, 1793–1797 (2014).

23. S. Sung, R. M. Amasino, Vernalization and epigenetics: how plants remember winter. Curr Opin Plant Biol 7, 4–10 (2004).

24. S. Berry, C. Dean, Environmental perception and epigenetic memory: mechanistic insight through FLC. Plant J 83, 133–148 (2015).

25. P. Crevillén, et al., Epigenetic reprogramming that prevents transgenerational inheritance of the vernalized state. Nature 515, 587–590 (2014).

26. E. J. Finnegan, M. Robertson, C. A. Helliwell, Resetting FLOWERING LOCUS C Expression After Vernalization Is Just Activation in the Early Embryo by a Different Name. Frontiers in Plant Science 11 (2021).

27. X. Luo, Y. He, Experiencing winter for spring flowering: A molecular epigenetic perspective on vernalization. J Integr Plant Biol 62, 104–117 (2020).

28. C. C. Sheldon, et al., Resetting of FLOWERING LOCUS C expression after epigenetic repression by vernalization. Proc Natl Acad Sci U S A 105, 2214–2219 (2008).

29. J. Choi, et al., Resetting and regulation of Flowering Locus C expression during Arabidopsis reproductive development. Plant J 57, 918–931 (2009).

30. Z. Tao, et al., Embryonic epigenetic reprogramming by a pioneer transcription factor in plants. Nature 551, 124–128 (2017).

31. Z. Tao, et al., Embryonic resetting of the parental vernalized state by two B3 domain transcription factors in Arabidopsis. Nat. Plants 5, 424–435 (2019).

32. G. Xu, Z. Tao, Y. He, Embryonic reactivation of FLOWERING LOCUS C by ABSCISIC ACID-INSENSITIVE 3 establishes the vernalization requirement in each Arabidopsis generation. Plant Cell 34, 2205–2221 (2022).

33. N. Sharma, et al., A Flowering Locus C Homolog Is a Vernalization-Regulated Repressor in Brachypodium and Is Cold Regulated in Wheat. Plant Physiol 173, 1301–1315 (2017).

34. A. Kennedy, K. Geuten, The Role of FLOWERING LOCUS C Relatives in Cereals. Front Plant Sci 11, 617340 (2020).

35. Q. Huan, Z. Mao, K. Chong, J. Zhang, Global analysis of H3K4me3/H3K27me3 in Brachypodium distachyon reveals VRN3 as critical epigenetic regulation point in vernalization and provides insights into epigenetic memory. New Phytologist 219, 1373–1387 (2018).

36. X. Liu, et al., Uncovering the transcriptional regulatory network involved in boosting wheat regeneration and transformation. Nat. Plants, 1–18 (2023).

37. L. Zhao, et al., Dynamic chromatin regulatory programs during embryogenesis of hexaploid wheat. Genome Biology 24, 7 (2023).

38. J. Guo, Y. Ren, Z. Tang, W. Shi, M. Zhou, Characterization and expression profiling of the ICE-CBF-COR genes in wheat. PeerJ 7, e8190 (2019).

39. W. Yong, et al., Vernalization-induced flowering in wheat is mediated by a lectin-like gene VER2. Planta 217, 261–270 (2003).

40. J. Ernst, M. Kellis, ChromHMM: automating chromatin-state discovery and characterization. Nat Methods 9, 215–216 (2012).

41. B. E. Bernstein, et al., A Bivalent Chromatin Structure Marks Key Developmental Genes in Embryonic Stem Cells. Cell 125, 315–326 (2006).

42. S. L. Klemm, Z. Shipony, W. J. Greenleaf, Chromatin accessibility and the regulatory epigenome. Nat Rev Genet 20, 207–220 (2019).

43. C. Li, et al., Wheat VRN1, FUL2 and FUL3 play critical and redundant roles in spikelet development and spike determinacy. Development 146, dev175398 (2019).

44. K. Wang, H. Liu, L. Du, X. Ye, Generation of marker-free transgenic hexaploid wheat via an Agrobacterium-mediated co-transformation strategy in commercial Chinese wheat varieties. Plant Biotechnology Journal 15, 614–623 (2017).

45. J. Xiao, D. Wagner, Polycomb repression in the regulation of growth and development in Arabidopsis. Current Opinion in Plant Biology 23, 15–24 (2015).

46. A. Pajoro, E. Severing, G. C. Angenent, R. G. H. Immink, Histone H3 lysine 36 methylation affects temperature-induced alternative splicing and flowering in plants. Genome Biol 18, 102 (2017).

47. N. Shitsukawa, et al., Wheat SOC1 functions independently of WAP1/VRN1, an integrator of vernalization and photoperiod flowering promotion pathways. Physiologia Plantarum 130, 627–636 (2007).

48. S. D. Michaels, R. M. Amasino, FLOWERING LOCUS C encodes a novel MADS domain protein that acts as a repressor of flowering. Plant Cell 11, 949–956 (1999).

49. N. Sharma, K. Geuten, B. S. Giri, A. Varma, The molecular mechanism of vernalization in Arabidopsis and cereals: role of Flowering Locus C and its homologs. Physiol Plant 170, 373–383 (2020).

50. M. Zikhali, L. U. Wingen, M. Leverington-Waite, S. Specel, S. Griffiths, The identification of new candidate genes Triticum aestivum FLOWERING LOCUS T3-B1 (TaFT3-B1) and TARGET OF EAT1 (TaTOE1-B1) controlling the short-day photoperiod response in bread wheat. Plant Cell Environ 40, 2678–2690 (2017).

51. T. S. Ream, D. P. Woods, R. M. Amasino, The molecular basis of vernalization in different plant groups. Cold Spring Harb Symp Quant Biol 77, 105–115 (2012).

52. S. Sung, R. M. Amasino, REMEMBERING WINTER: Toward a Molecular Understanding of Vernalization. Annual Review of Plant Biology 56, 491–508 (2005).

53. J. Song, A. Angel, M. Howard, C. Dean, Vernalization – a cold-induced epigenetic switch. Journal of Cell Science 125, 3723–3731 (2012).

54. P. Ruelens, et al., FLOWERING LOCUS C in monocots and the tandem origin of angiosperm-specific MADS-box genes. Nat Commun 4, 2280 (2013).

55. International Wheat Genome Sequencing Consortium (IWGSC), Shifting the limits in wheat research and breeding using a fully annotated reference genome. Science 361, eaar7191 (2018).

56. A. G. Greenup, et al., ODDSOC2 Is a MADS Box Floral Repressor That Is Down-Regulated by Vernalization in Temperate Cereals. Plant Physiology 153, 1062–1073 (2010).

57. C. B. Field, V. Barros, T. F. Stocker, Q. Dahe, Eds., Managing the Risks of Extreme Events and Disasters to Advance Climate Change Adaptation: Special Report of the Intergovernmental Panel on Climate Change (Cambridge University Press, 2012) 10.1017/CBO9781139177245 (December 19, 2023).

58. D. Wang, et al., Boosting wheat functional genomics via an indexed EMS mutant library of KN9204. Plant Comm 4 (2023).

59. S. Chen, Y. Zhou, Y. Chen, J. Gu, fastp: an ultra-fast all-in-one FASTQ preprocessor. Bioinformatics 34, i884–i890 (2018).

60. D. Kim, J. M. Paggi, C. Park, C. Bennett, S. L. Salzberg, Graph-based genome alignment and genotyping with HISAT2 and HISAT-genotype. Nat Biotechnol 37, 907–915 (2019).

61. P. Danecek, et al., Twelve years of SAMtools and BCFtools. Gigascience 10, giab008 (2021).

62. Y. Liao, G. K. Smyth, W. Shi, featureCounts: an efficient general purpose program for assigning sequence reads to genomic features. Bioinformatics 30, 923–930 (2014).

63. M. I. Love, W. Huber, S. Anders, Moderated estimation of fold change and dispersion for RNA-seq data with DESeq2. Genome Biol 15, 550 (2014).

64. Z. Gu, R. Eils, M. Schlesner, Complex heatmaps reveal patterns and correlations in multidimensional genomic data. Bioinformatics 32, 2847–2849 (2016).

65. T. Wu, et al., clusterProfiler 4.0: A universal enrichment tool for interpreting omics data. Innovation (Camb) 2, 100141 (2021).

66. H. Mi, A. Muruganujan, D. Ebert, X. Huang, P. D. Thomas, PANTHER version 14: more genomes, a new PANTHER GO-slim and improvements in enrichment analysis tools. Nucleic Acids Res 47, D419–D426 (2019).

67. F. Tian, D.-C. Yang, Y.-Q. Meng, J. Jin, G. Gao, PlantRegMap: charting functional regulatory maps in plants. Nucleic Acids Res 48, D1104–D1113 (2020).

68. H. Li, R. Durbin, Fast and accurate short read alignment with Burrows-Wheeler transform. Bioinformatics 25, 1754–1760 (2009).

69. F. Ramírez, et al., deepTools2: a next generation web server for deep-sequencing data analysis. Nucleic Acids Res 44, W160–165 (2016).

70. H. Thorvaldsdóttir, J. T. Robinson, J. P. Mesirov, Integrative Genomics Viewer (IGV): high-performance genomics data visualization and exploration. Brief Bioinform 14, 178–192 (2013).

71. Y. Zhang, et al., Model-based analysis of ChIP-Seq (MACS). Genome Biol 9, R137 (2008).

72. A. R. Quinlan, I. M. Hall, BEDTools: a flexible suite of utilities for comparing genomic features. Bioinformatics 26, 841–842 (2010).

73. G. Yu, L.-G. Wang, Q.-Y. He, ChIPseeker: an R/Bioconductor package for ChIP peak annotation, comparison and visualization. Bioinformatics 31, 2382–2383 (2015).

74. S. Cheng, et al., Harnessing Landrace Diversity Empowers Wheat Breeding for Climate Resilience. 2023.10.04.560903 (2023).

75. L. S. Johnson, S. R. Eddy, E. Portugaly, Hidden Markov model speed heuristic and iterative HMM search procedure. BMC Bioinformatics 11, 431 (2010).

76. R. C. Edgar, Muscle5: High-accuracy alignment ensembles enable unbiased assessments of sequence homology and phylogeny. Nat Commun 13, 6968 (2022).

77. S. Capella-Gutiérrez, J. M. Silla-Martínez, T. Gabaldón, trimAl: a tool for automated alignment trimming in large-scale phylogenetic analyses. Bioinformatics 25, 1972–1973 (2009).

78. A. Stamatakis, RAxML version 8: a tool for phylogenetic analysis and post-analysis of large phylogenies. Bioinformatics 30, 1312–1313 (2014).

79. Y. Wang, et al., Simultaneous editing of three homoeoalleles in hexaploid bread wheat confers heritable resistance to powdery mildew. Nat Biotechnol 32, 947–951 (2014).

80. Q. Shan, et al., Targeted genome modification of crop plants using a CRISPR-Cas system. Nat Biotechnol 31, 686–688 (2013).

81. R. Cui, et al., Functional conservation and diversification of class E floral homeotic genes in rice (Oryza sativa). Plant J 61, 767–781 (2010).

82. K. J. Livak, T. D. Schmittgen, Analysis of relative gene expression data using real-time quantitative PCR and the 2(-Delta Delta C(T)) Method. Methods 25, 402–408 (2001).

83. C. Tian, et al., Impaired histone inheritance promotes tumor progression. Nat Commun 14, 3429 (2023).

84. X. Lin, et al., Systematic mining and functional analysis of factors regulating wheat spike development for breeding selection. 2022.11.11.516122 (2023).

85. S. Schilling, A. Kennedy, S. Pan, L. S. Jermiin, R. Melzer, Genome-wide analysis of MIKC-type MADS-box genes in wheat: pervasive duplications, functional conservation and putative neofunctionalization. New Phytol 225, 511–529 (2020).

